# Rapid ex vivo reverse genetics identifies the essential determinants of prion protein toxicity

**DOI:** 10.1101/2022.10.05.509883

**Authors:** Regina R. Reimann, Martina Puzio, Antonella Rosati, Marc Emmenegger, Bernard L Schneider, Pamela Valdés, Danzhi Huang, Amedeo Caflisch, Adriano Aguzzi

**Affiliations:** Institute of Neuropathology, University of Zurich, Schmelzbergstrasse 12, CH-8091 Zurich, Switzerland; Brain Mind Institute, Ecole Polytechnique Fédérale de Lausanne, Station 19, 1015 Lausanne, Switzerland; Bertarelli Foundation Gene Therapy Platform, Ecole Polytechnique Fédérale de Lausanne, Ch. des Mines 9, 1202 Geneva, Switzerland; Department of Biochemistry, University of Zürich, CH-8057 Zürich, Switzerland

## Abstract

The cellular prion protein PrP^C^ mediates the neurotoxicity of prions and other protein aggregates through poorly understood mechanisms. Antibody-derived ligands against the globular domain of PrP^C^ (GDL) can also initiate neurotoxicity by inducing an intramolecular R_208_-H_140_ hydrogen bond (“H-latch”) between the α2-α3 and β2-α2 loops of PrP^C^. Importantly, GDL that suppress the H-latch prolong the life of prion-infected mice, suggesting that GDL toxicity and prion infections exploit convergent pathways. To define the structural underpinnings of these phenomena, we transduced nineteen individual PrP^C^ variants to PrP^C^-deficient cerebellar organotypic cultured slices using adenovirus-associated viral vectors (AAV). We report that GDL toxicity requires a single N-proximal cationic residue (K_27_ or R_27_) within PrP^C^. Alanine substitution of K_27_ also prevented the toxicity of PrP^C^ mutants that induce Shmerling syndrome, a neurodegenerative disease that is suppressed by co-expression of wild-type PrP^C^. K_27_ may represent an actionable target for compounds aimed at preventing prion-related neurodegeneration.

## Introduction

The structure of the cellular prion protein PrP^C^ encompasses an amino-proximal flexible tail (FT, amino acid residues 23-123) linked to a globular domain (GD, 124-230) [1]. PrP^C^ triggers a neurotoxic cascade upon interaction with prions [2-4], other pathological protein aggregates including Aβ [5-7], and antibodies or antibody-derived ligands against its globular domain (henceforth collectively called globular-domain ligands or GDL) [8-11]. In addition, PrP^C^ mutants carrying deletions of certain domains induce severe neurodegeneration, which can be suppressed by co-expression of wild-type (wt) PrP^C^ [12-14].

We and others have found that ligands targeting the α1 and α3 helices of the GD, or the second charge cluster (CC2: 95-110), induce an acute neurodegenerative phenotype [9-11] resembling that of prion infections [15]. This toxic function of GDL requires the induction of an intramolecular R28-H140 hydrogen bond [16]. Because their toxicity is rapid and the structural details of their interaction with PrP^C^ are atomistically understood, these ligands represent useful tools in the investigation of prion-initiated neurodegeneration [17], and have enabled the identification of the FT as the mediator of neurotoxicity [9].

The FT can be dissected into distinct domains: a polybasic region (PR: residues 23-31) encompassing an N-proximal charge cluster (CC1: residues 23-27), a region preceding the octapeptide repeat region (pre-OR: residues 28-49), the five octapeptide repeats (OR: residues 50-90), a second charge cluster (CC2) and a hydrophobic core (HC: residues 111-134) [18]. These structural domains play distinct roles in PrP^C^-mediated neurotoxicity, as shown by the differential patterns of prion-associated degeneration in mice expressing the cognate deletion mutants of PrP^C^ [18].

Here we have perturbed the sequence of the FT in order to delineate the requirements for neurotoxicity. Using AAV-mediated expression of NG and various PrP^C^ mutants in cerebellar organotypic slice cultures (COCS), we have assessed the toxicity of nineteen individual PrP^C^ mutants (Table 1 and Figure 1A). We identified a lysine residue at position 27 of the *Prnp* reading frame as essential for the neurodegeneration induced by globular domain ligands (GDL). Furthermore, we found that co-expression of a PrP^C^ variant with an uncharged CC1suppresses the neurodegeneration induced by toxic PrP^C^ from which the HC was deleted.

**Table 1:**
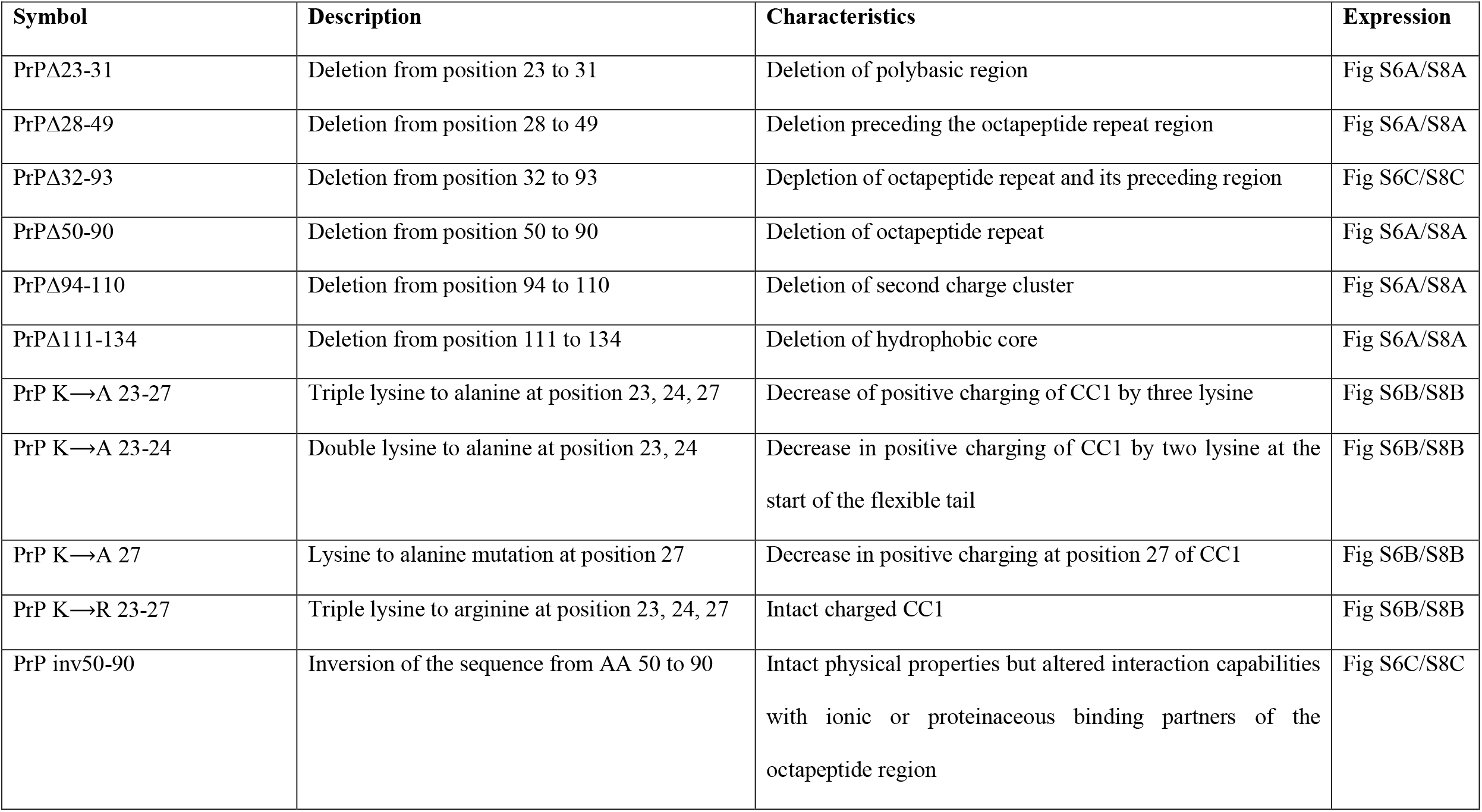
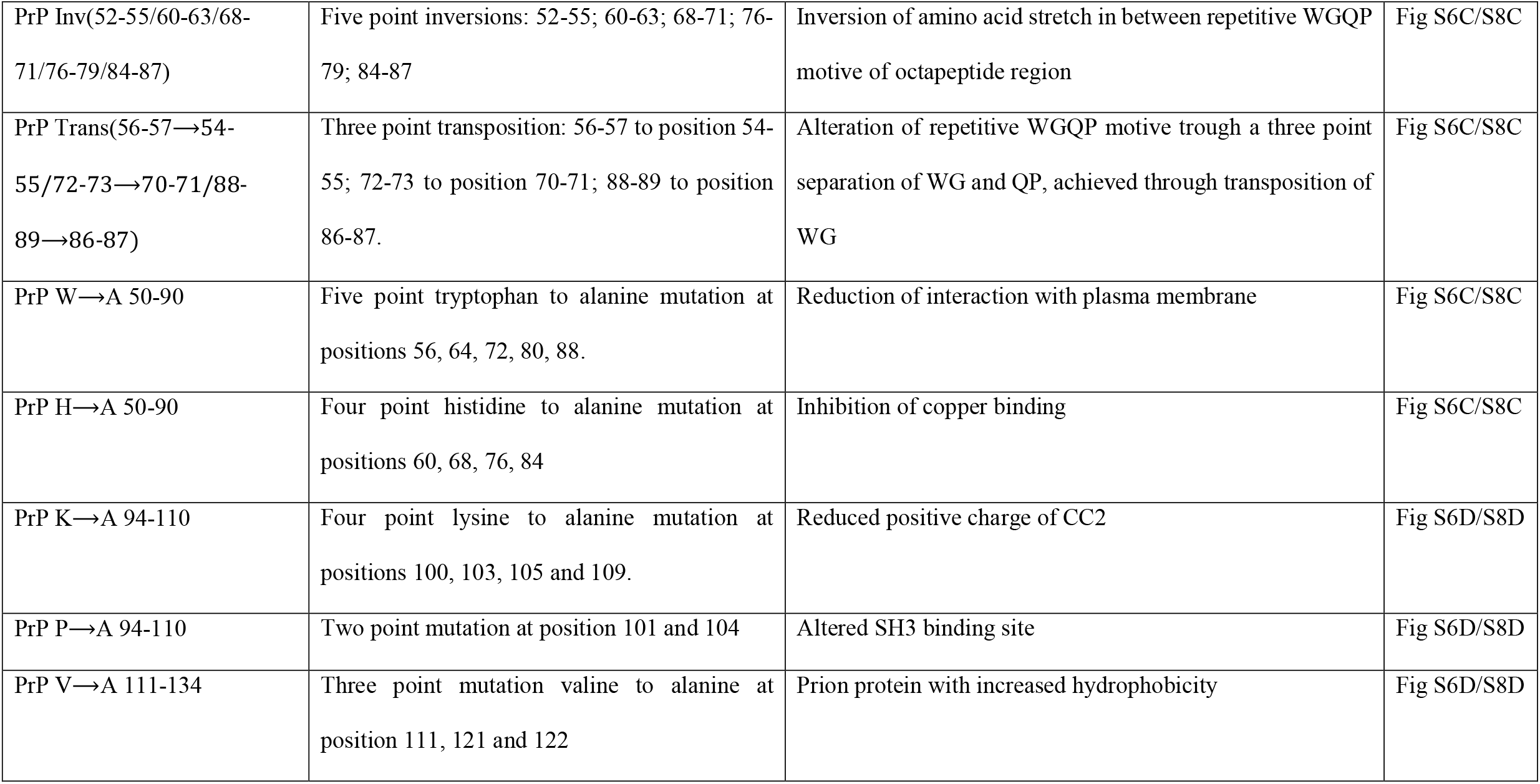
Summary prion protein constructs.

**Figure 1:**
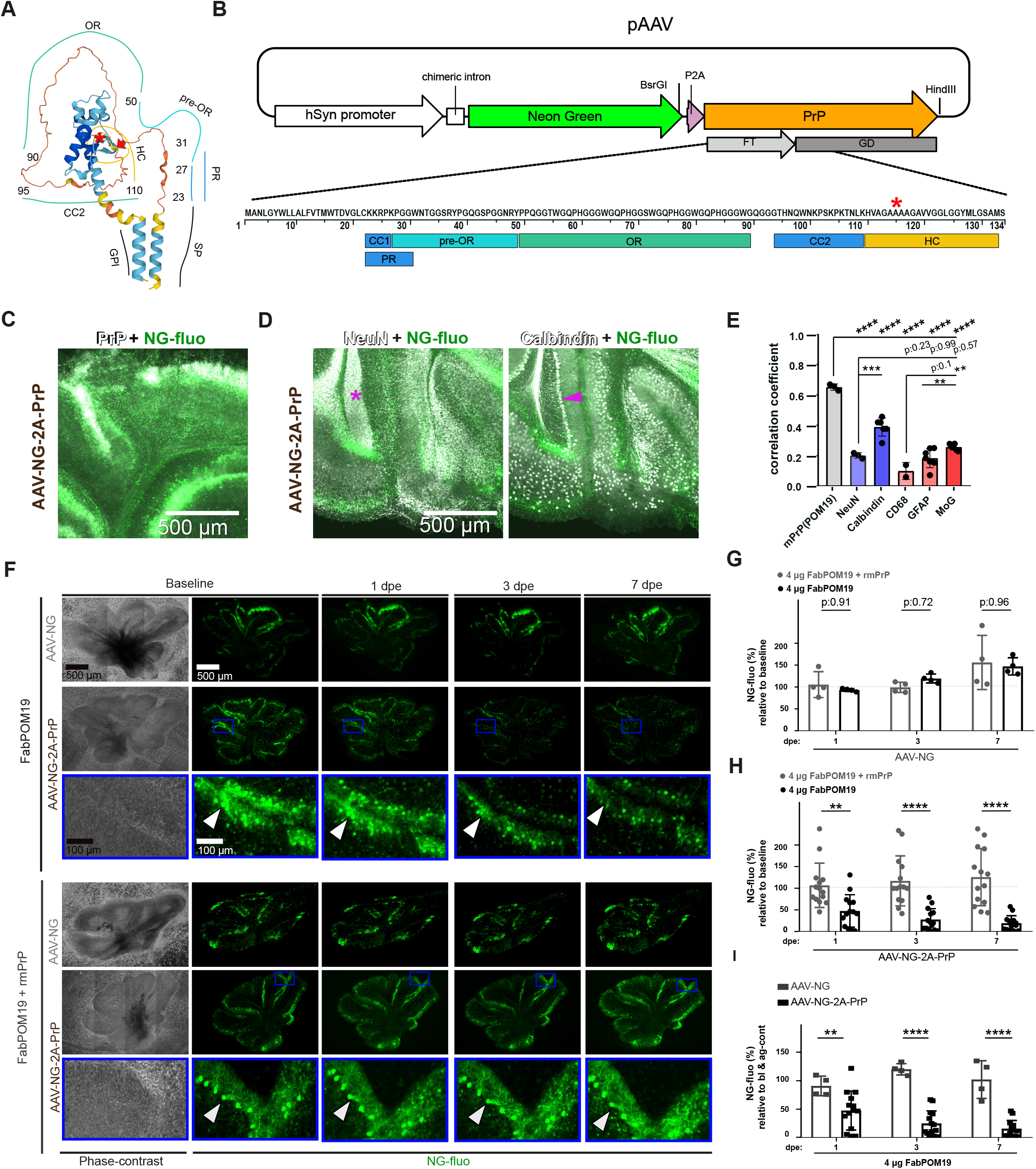
GDL-induced Purkinje cell degeneration in Prnp^ZH3/ZH3^ COCS transduced with PrP^-C^. (**A**) Structure of the full-length murine prion protein, as visualized by AlphaFold [57, 58]. The numbers indicate residue positions. SP: signaling peptide. GPI: glycosylphosphatidylinositol anchor. CC1: charge cluster 1. PR: polybasic region. Pre-OR: region preceding the octapeptide repeats (OR). CC2: charge cluster 2. HC: hydrophobic core. The red asterisk denotes the border between the flexible tail (FT) and the globular domain (GD) at residue 123/124. Red arrowhead: residue 134. (**B**) Schema of the polyprotein vector. A P2A self-cleavage site enables the coordinated expression of NeonGreen (NG) and the murine wt or mutated prion protein (PrP) driven by the human synapsin 1 (hSyn1) promoter. The replacement of wt *Prnp* with cassettes encoding mutated *Prnp* versions was performed using the BsrGI and HindIII restriction sites. FT: light grey. GD: dark grey. Red asterisk: border between FT and GD. Colored blocks illustrate the different domains of the FT. Light blue: CC1 and CC2; turquoise: pre-OR; green: OR; orange: HC. (**C-D**) Representative micrographs of cerebellar organotypic slice cultures (COCS) 19 days post transduction with AAV-hSyn1-NG-2A-PrP. (**C**) Immunofluorescence staining for PrP with POM19 (white) demonstrates a high correlation of expression. Green: fluorescent signal for Neon Green. (**D**) Co-immunostaining for neuronal protein markers localize NG-fluorescence expression predominantly to Purkinje cells (magenta arrowhead) rather than to granule cells (magenta asterisk). Neuronal nuclear antigen (NeuN, left panel, in white) expressed in cerebellar granule cells and Calbindin (right panel, white) expressed in Purkinje cells. Green fluorescence signal: NeonGreen. (**E**) Immunofluorescence colocalisation analysis (Pearson’s coefficient) of different markers with NG-fluo. Representative images of this analysis are shown in C, D and in S1 B-E. The NeonGreen signal is highly correlated with POM19 staining (PrP) and with calbindin. Blue bars: Neuronal markers (NeuN and calbindin). Red bars: non-neuronal markers: CD68 (microglia), glial fibrillary acidic protein (GFAP, expressed by astrocytes) and myelin-oligodendrocyte glycoprotein (MoG, expressed by oligodendrocytes). Each dot represents one COCS, mean±sd; **: p<0.01; ***: p<0.001; ****: p<0.0001. Two-way Anova, Tukey post hoc test. (**F**) Phase-contrast (left) and fluorescence micrographs (right) of *Prnp*^ZH3/ZH3^ COCS two weeks after transduction with AAV-hSyn1-NG-2A-PrP or AAV-hSyn1-NG (10^9^ TU/ml, leftmost column), and at various time points after a pulse of FabPOM19 (4 μg in 10 μl PBS). NG expression by Purkinje cells was initially conspicuous but became gradually reduced under FabPOM19 treatment, and was almost completely lost after one week. Pre-incubation of FabPOM19 with a 3x molar excess of PrP^C^ (lower three rows) prevented the disappearance of NG fluorescence. Blue rectangles: magnified regions. White arrowhead: bodies of individual Purkinje cells. (**G**) NeonGreen (NG) morphometric analysis (details: Fig. S4) revealed no significant degeneration of NG-only expressing COCS treated with FabPOM19. Data points: percentage of NG expression at baseline. AAV titer: 10^9^ TU/ml; each dot represent one COCS, mean ± sd, two-Way Anova, Sidak post hoc test. Dpe: Days post exposure to FabPOM19 or blocked control. (**H**) Progressive, massive decrease in NG expression over time (24 hours, 72 hours, 1 week) in slices infected with AAV-Syn-NG-2A-PrP and subjected to FabPOM19 treatment. AAV titer: 10^9^ TU/ml. Each dot represent one COCS. Mean ±sd, two-way anova, Sidak post hoc test; ns=not significant, ****P<0.0001, two-way anova, Sidak post hoc test. Dpe: Days post exposure. (**I**) Data points from G and H additionally divided by the average NG expression of COCS treated with blocked antibody at the same time point. The data representation used in this figure for the neurodegeneration assay is also used in the following figures. Each dot represents one COCS, mean±sd, ns=not significant, *P < 0.05, **P<0.01, *+***P<0.0001 one-Way Anova, Sidak post hoc test. Dpe: Days post exposure.

## Results

### An ex vivo assay for probing domains of PrP relevant to neurodegeneration

To investigate the role of the different domains of the FT of PrP^C^ in neurotoxicity, we measured the damage to COCS expressing various PrP variants after exposure to GDLs. To reliably assess the progressive disappearance of cells in organotypic slices, we established a system for simultaneous transduction of PrP^C^ mutants alone with the genetically encoded fluorescent marker, Neon Green (NG). We reasoned that the progressive disappearance of NG fluorescence over time would point to the degeneration of the respective cells. We thus assembled a polyprotein gene consisting of various mutants of the murine *Prnp* gene (encoding PrP^C^) followed by a P2A self-cleaving sequence and NG. This transgenic cassette was placed under the transcriptional control of the neuron-specific human synapsin 1 (hSyn1) promoter (**Fig 1B and S1 A**). We expressed this construct transiently using adeno-associated viral AAV2/6 (AAV6 capsid with AAV2 inverted terminal repeats) vectors in cerebellar organotypic cultured slices (COCS) from PrP-deficient mice (*Prnp*^ZH3/ZH3^, [19]) and found conspicuous NG expression in Purkinje cells as expected from this particular AAV serotype (**Fig 1 C-E and S1 B-E**).

Since mice lacking PrP^C^ are resistant to GDL-mediated neurotoxicity [9], they provide an ideal platform for dissecting the functional domains of PrP^C^ by reverse genetics. We exposed transfected *Prnp*^ZH3/ZH3^ COCS (NG versus NG-2A-PrP) to chronic anti-PrP antibody treatment [9]. After 10 days of exposure to 134 nM of Fab-POM19 (which recognizes a discontinuous PrP epitope comprising residues 121-134 and 218-221 of the GD) in culture medium (5 media changes corresponding to a cumulative dose of 3.7 μg/COCS), we found that transduction restored GDL-mediated neurodegeneration of *Prnp*^ZH3/ZH3^ COCS. Conversely, slices transduced with an empty AAV vector (AAV-NG) were resistant to GDLs (**Fig S2A-D**).

To accelerate, challenge and validate our experimental system, we added 4 μg of the GDL Fab-POM19 (which recognizes a discontinuous PrP epitope comprising residues 121-134 and 218-221) in the form of directly trickled drops (henceforth called “pulse treatment”) on *Prnp*-overexpressing *tg*a*20* COCS and performed several analyses at various time points after exposure. As expected, we observed a severe loss of Neuronal nuclear antigen (NeuN, **Fig S3A-D**) within 24 hours both by Western blotting as well as increased propidium iodide (PI) staining, a marker of cell-membrane damage (**Fig S3E-G**).

In order to optimize the imaging and quantification of NG expression, we cultured the slices directly on coverslips which were placed into roller tubes [20]. We transduced PrP^C^ into COCS using AAV vectors (AAV-NG-2A-PrPwt), performed baseline confocal imaging (19 days post transduction) at 5 different depths, and generated maximal-intensity projections (**Fig S4A-B**) followed by NG morphometry (**Fig S4 C-E**). With this method NG expression was stable for ≥4 weeks post transduction (**Fig S4 F-G**). Immediately after baseline scanning, we pulse-treated each COCS with 4 μg of FabPOM19 or, for control, FabPOM19 inactivated by preincubation with its cognate antigen. Then, we reimaged COCS in a time-course experiment at 1, 3 and 7 days post exposure (dpe) to treatment using identical confocal imaging setting as for baseline imaging, and compared NG expression of each COCS and at each time-point with its baseline expression. Loss of NG expression, indicating incipient neurodegeneration, was found already at 1 dpe, with increasing severity at 3 and 7 dpe (**Fig 1F-I**), and a trend towards increased PI incorporation by NG-expressing Purkinje cells (*p* = 0.1), corroborating the suggestion that NG loss was the result of cell death (**Fig S5A-B**). We obtained similar results when repeating this experiment with a different neurotoxic GDL (FabPOM1; binding residues: 138-147; 204; 208; 212) (**Fig S5 C**).

### Role of the octarepeats in anti-PrP antibody mediated neurodegeneration

Mice overexpressing PrP_Δ32-93_ are resistant to toxic GDLs [9]. Accordingly, we found no significant reduction of NG expression (p= 0.42 at 7 days post exposure) when COCS were treated with toxic GDLs two weeks after transduction with AAV-NG-2A-PrP_Δ32-93_ (**Fig 2A, Table 1**) despite intact surface expression of the mutant protein (**Fig S6C**). The FT may exert its toxic function via a pathological interaction with membrane constituents [21]. The OR features a high tryptophan content which may favor its insertion into biological membranes [22], suggesting that the FT may exert its pathological function by integrating the OR into the plasma membrane. We examined this possibility using molecular dynamics (MD) simulations (**Fig S7A**). We found that a potential OR insertion into the membrane can occur spontaneously within a 100-ns time scale simulated for the PrP60-83 peptide (**Fig S7B)**, respectively within 50-ns for the PrP21-90 peptide (**Fig S7C**). Furthermore, a three-time tryptophan-to-alanine mutation in the PrP60-83 peptide would block this interaction (**Fig 2B, Fig S7D**). In order to test this observation experimentally, we transduced into *Prnp*^ZH3/ZH3^ COCS a *Prnp* mutant in which all tryptophan residues in the OR region were replaced with alanines (**Table 1**). The protein expression levels of the mutant were similar to those of wt PrP^C^ (**Fig S9A**). This manipulation was expected to prevent any insertion events (**Fig 2B**). However, we found that the tryptophan-mutated construct restored GDL-mediated neurodegeneration (**Fig 2C**). This makes the membrane-insertion hypothesis of GDL-mediated neurodegeneration unlikely. Recent reports suggest that the neurotoxic activity of the FT is inhibited by a copper-mediated *cis* interaction with the GD [23, 24]. If so, the four histidine residues in the OR, which bind Cu^2+^ with sub-nanomolar affinity [25], would be essential for this phenomenon. To test the influence of these residues on GD-ligand mediated neurodegeneration, we replaced all four histidine in the OR with alanine within the OR (**Table 1**). However, we did not detect any reduction of GDL-induced neurodegeneration in comparison to wt PrP (**Fig 2C**). The distance between the two charge clusters in the PrP_Δ32-93_ deletion mutants is reduced, with possible secondary effects on protein function that may not be directly linked to the OR. We generated a construct with an entirely inverted OR (**Table 1**), which is expected to show similar physical properties as wt PrP^C^ but altered interaction capabilities with ionic or proteinaceous binding partners. Protein surface expression and glycosylation of the mutant was intact (**Fig S6C, Fig S8C**). Interestingly, this construct did not restore GDL-induced neurodegeneration (**Fig 2D**). In an attempt to narrow down the critical amino acid motifs in the OR down more precisely, we designed two additional PrP^C^ mutants based on the same concept but with smaller modifications. On one hand, we generated a construct in which the repetitive WGQP motifs are intact but the stretches in-between are inverted and on the other hand, we modified exactly this motif at three positions by translocation of WQ (**Table 1**). As both mutations restored GDL-mediated neurodegeneration (**Fig 2E**), this approach was not successful in the further identification of OR amino-acids motifs important for mediating neurodegeneration. Since GDL-mediated neurodegeneration is dependent on PrP^C^ expression level [9], we asked whether PrP_Δ32-93_ might mediate neurodegeneration if overexpressed even stronger in Purkinje cells. Indeed, at a threefold higher expression level than assessed previously (**Fig S9A**), PrP_Δ32-93_ restored GDL-mediated neurodegeneration (**Fig 2F**). In contrast to wt PrP expressed at a comparably high level, the induction of degeneration was delayed by one day. This finding suggests that the OR is an important, but not essential, modulator of anti-prion GDL-mediated neurodegeneration.

**Figure 2:**
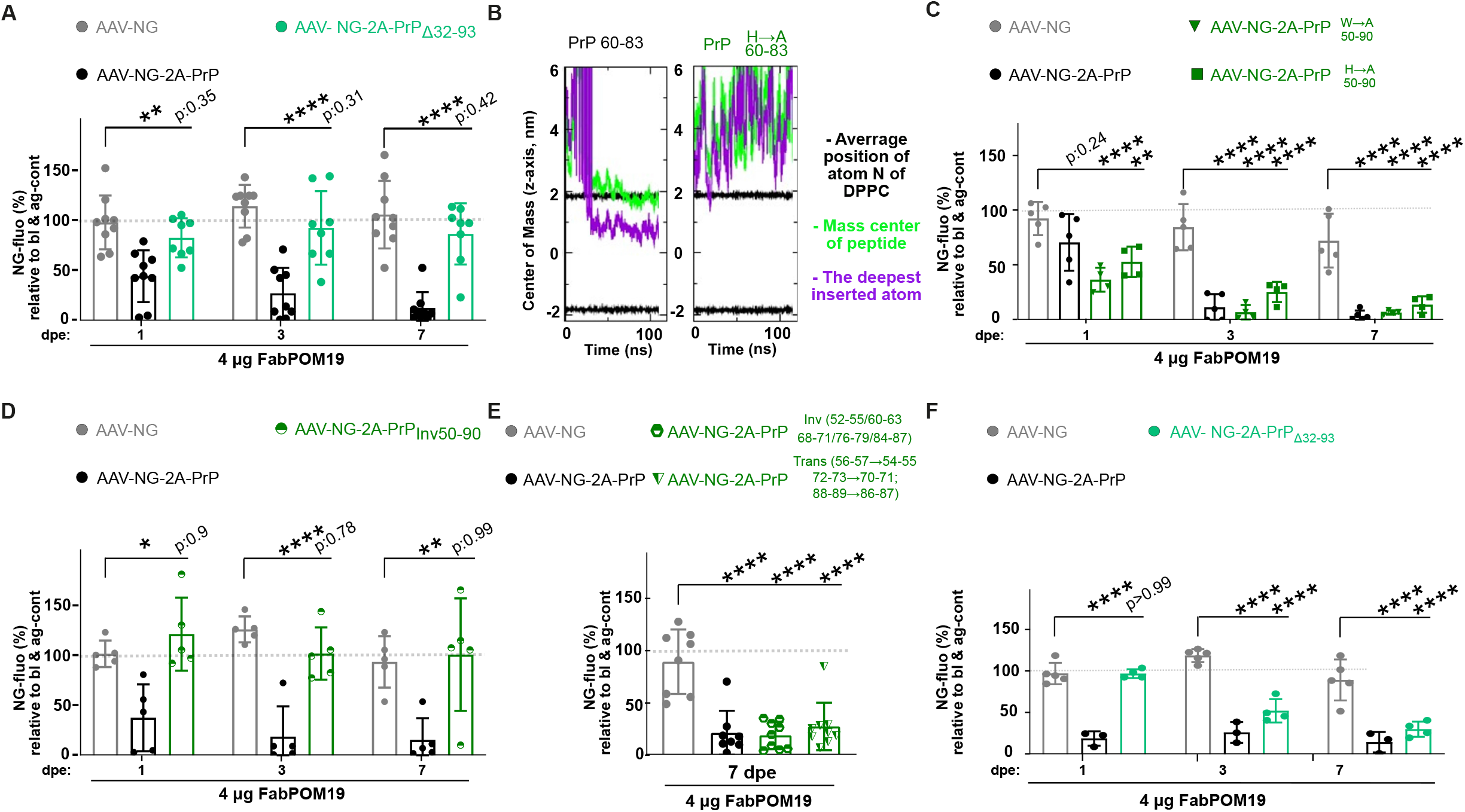
The OR modulates GDL-mediated neurodegeneration but is not essential. (**A**) Morphometric NG time-course analysis showing that PrP^C^ but not PrP_Δ32-93_ restore GDL-induced neurodegeneration in *Prnp*^ZH3/ZH3^ COCS. AAV titer: 1 E+9 TU/ml. Here and in all following panels if not otherwise specified: Each dot represents one COCS, *P<0.05, **P<0.01, ***P<0.001, ****P<0.0001; each data point in percentage of NG expression at baseline is divided by the average NG expression of COCS treated with blocked antibody at the same time point; one-Way anova, Sidak post hoc test. Dpe: days post exposure. (**B**) Molecular dynamics simulation of the PrP60-83 peptide with a three-point tryptophan to alanine mutant (right) showed no insertion of the FT into the plasma membrane in contrast to wt PrP60-83 (left). Further details on this experiment are given in Fig S7. (**C-D**) Time course analysis demonstrates full restoration of neurodegeneration involving 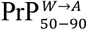 and 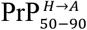 mutants (C), but not PrP_Inv(50-90)_ with an inversed OR sequence (D). (**E**) Neither a flip of components of the 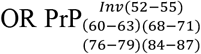 nor a three-point transposition of tryptophan-glycine elements 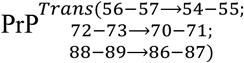 blocks FabPOM19 mediated neurodegeneration. One-way anova, Dunnett’s post hoc test. (**F**) Repetition of the experiment performed in B, but with COCS exposed to a 10-fold higher AAV titer, resulting in approximatively 3-fold higher expression (**Fig S9A**) and faster kinetic of NG loss. COCS expressing PrP_Δ32-93_ remained intact after one day of FabPOM19 exposure, but experienced severe neurodegeneration at later time points.

### A lysine residue at position 27 is essential for PrP^C^ mediated neurodegeneration

We then systematically analyzed various mutants of PrP^C^ carrying deletions in each of its functional units (**Table 1**). To robustly identify the most relevant regions for GD mediated neurodegeneration, COCS were infected with a five-fold higher functional titer than used in our previous experiments. This resulted in two-fold higher PrP expression level (**Fig S9B**). We found that all constructs, except those lacking the PR (PrP_Δ23-31_), were able to restore GDL-mediated neurodegeneration (**Fig 3A**). However, all mutants showed a one-day delay in the velocity of induction. In addition, PrP_Δ111-134_ showed low expression in CAD5 cells (**Fig S8A**) and no expression in COCS (**Fig S9B**), probably recapitulating the previously described phenotype of mice expressing deletion mutants of PrP in this region [26]. We excluded this construct from further analyses.

**Figure 3:**
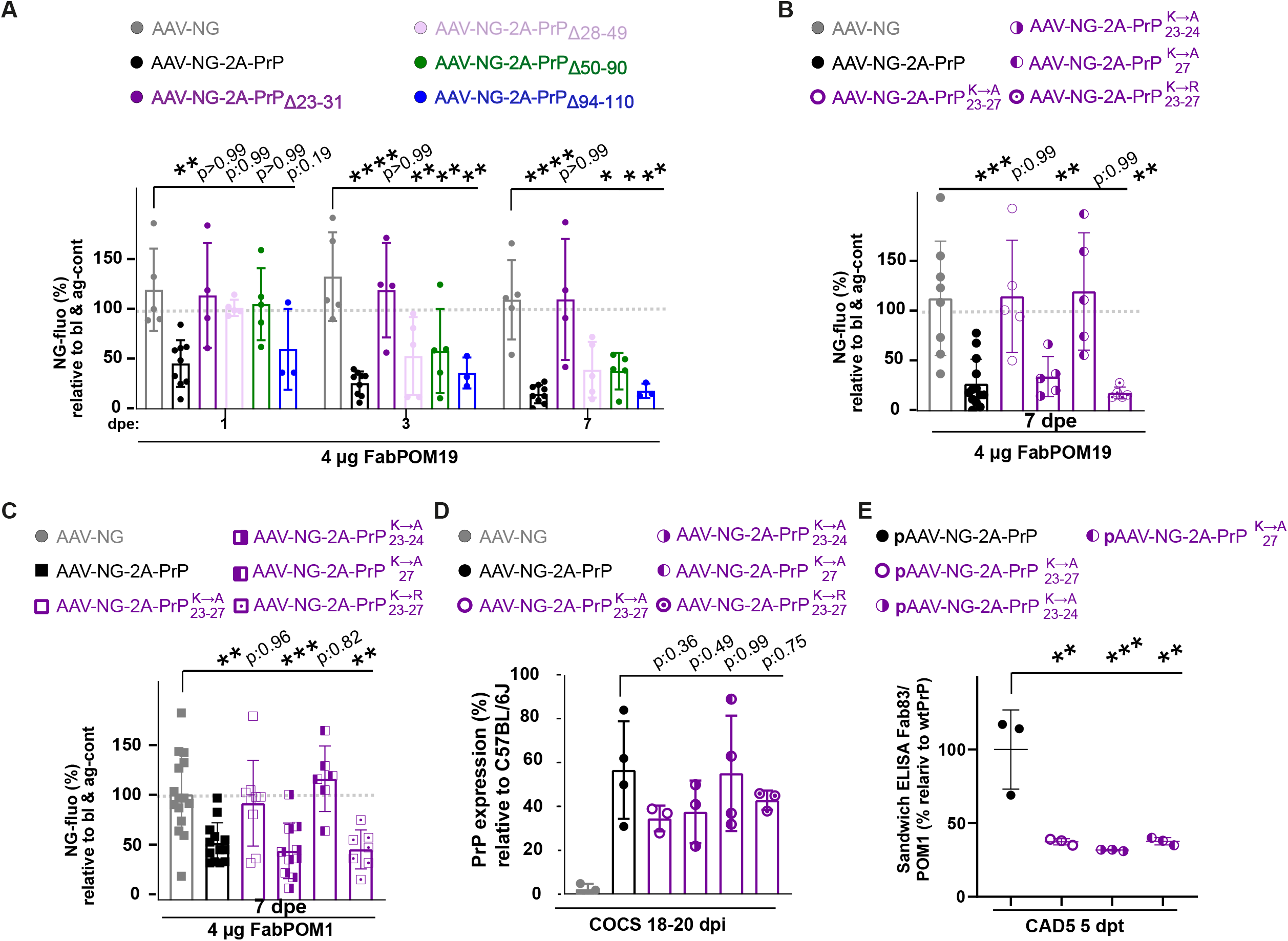
A LYS_27_->ALA point mutation suppresses GDL-mediated neurodegeneration. (**A**) Time-dependent FabPOM19-induced neurodegeneration in COCS expressing various *Prnp* deletion mutants. All mutants except those with a deletion of the polybasic region (violet) restored FabPOM19 toxicity. Here and henceforth: AAV titer 5 E+9 TU/ml, each dot represent one COCS, *P<0.05, **P<0.01, ***P<0.001, ****P<0.0001. One-Way Anova, Sidak post hoc test. Datapoints: percentage of NG expression at baseline divided by the average NG expression of COCS treated with paratope-blocked antibody at the same time point. (**B**) Prion protein mutants with all K->A mutations within the polybasic region, or with K->A only at position 27, do not restore anti-prion antibody-mediated neurodegeneration induced by FabPOM19. In contrast, replacing the lysines at position 23 and 24 with alanines did not modify FabPOM19 toxicity. K->R point mutations had no impact on FabPOM19 toxicity. (**C**) The experiment shown in B was replicated with the toxic GDL FabPOM1. (**D**) PrP^C^ expression level from COCS transfected together with those used for the experiments in B and C but grown separately on an inlay demonstrate similar expression levels of wt versus CC1 mutant PrP. Expression levels shown are relative to PrP^C^ expression levels of cerebellar homogenate from C57BL/6J wt mice. Sandwich ELISA (coating: POM1; probing biotinylated POM19). One dot represents one inlay with 11 slices. (**E**) ELISA (Coating with Fab83, incubating with wt or altered PrP^C^, and probing with biotinylated FabPOM1) assessing the binding capacity of Fab83 to the three CC1-mutated PrP^C^ versions. Mutation of K23 and K24 had a similar effect as the single-point mutation of K27. All concentrations relative to wtPrP^C^ 1 five days after transduction of CAD5 cells with pAAV vectors. ELISA data were performed in triplicates. ** p<0.01, *** p<0.001.

Electrophysiological assays in cells had established the importance of the PR as the main effector of GDL-mediated (POM1, POM11, D18) pathological inward current, as well as the importance of positively charged amino acids within this domain for triggering spontaneous currents in cells expressing PrP^C^ with a deletion in the central domain (PrP_Δ105-125_) [11]. We therefore asked whether these charged amino acids are equally important for GDL-mediated neurodegeneration in COCS. We mutated the three lysine residues of the CC1 to alanine (**Table 1**). As with to the polycationic region-depleted mutant, PrP_*K*→*A* 23−27_ did not restore anti-prion antibody-mediated neurodegeneration (**Fig 3B-C**). In order to establish whether all lysine residues are equally important, we compared a construct with lysine-to-alanine conversions of the first two N-terminal lysines (24, 25) to conversion of lysine at position 27 (**Table 1**). Surprisingly, the point mutation of lysine 27 proved sufficient to block GDL toxicity, probed with FabPOM1 and FabPOM19 **(Fig 3B-C**), whereas PrP_*K*→*A* 23−24_ fully rescued GDL-mediated neurodegeneration. Protein expression levels were similar for all constructs (**Fig 3D**). We then asked whether the effect is charge-dependent by expressing a PrP_*K*→*R* 23−27_ construct (**Table 1**). This construct restored GDL-mediated neurodegeneration (**Fig 3B-C**). In line with these data, we found that the CC1 ligand Fab83, which interacts with the tree N-terminal lysines, is neuroprotective in prion-infected COCS [27]. We therefore used ELISA to investigate the binding capacity of Fab83 to the protein mutants PrP_*K*→*A* 23−24_ versus PrP_*K*→*A*27_ and found a similar binding reduction (**Fig 3E**).

### CC2 ligand neurotoxicity is CC1 dependent

Analogously to previous studies [9], deletion of CC2 had no effect on GDL-mediated neurodegeneration (**Fig 3A**). Moreover, PrP^C^ mutants with a deionized CC2 or with an altered Src homology 3 (SH3) [28] domain restored GDL-mediated neurodegeneration (**Fig 4A**). Several studies have reported a high-affinity interaction of the CC2 with amyloid-β (Aβ) oligomers [5, 29]. Additionally, PrP^C^ may act as a receptor for Aβ-oligomers mediating impairment of synaptic plasticity [5, 6].

**Figure 4:**
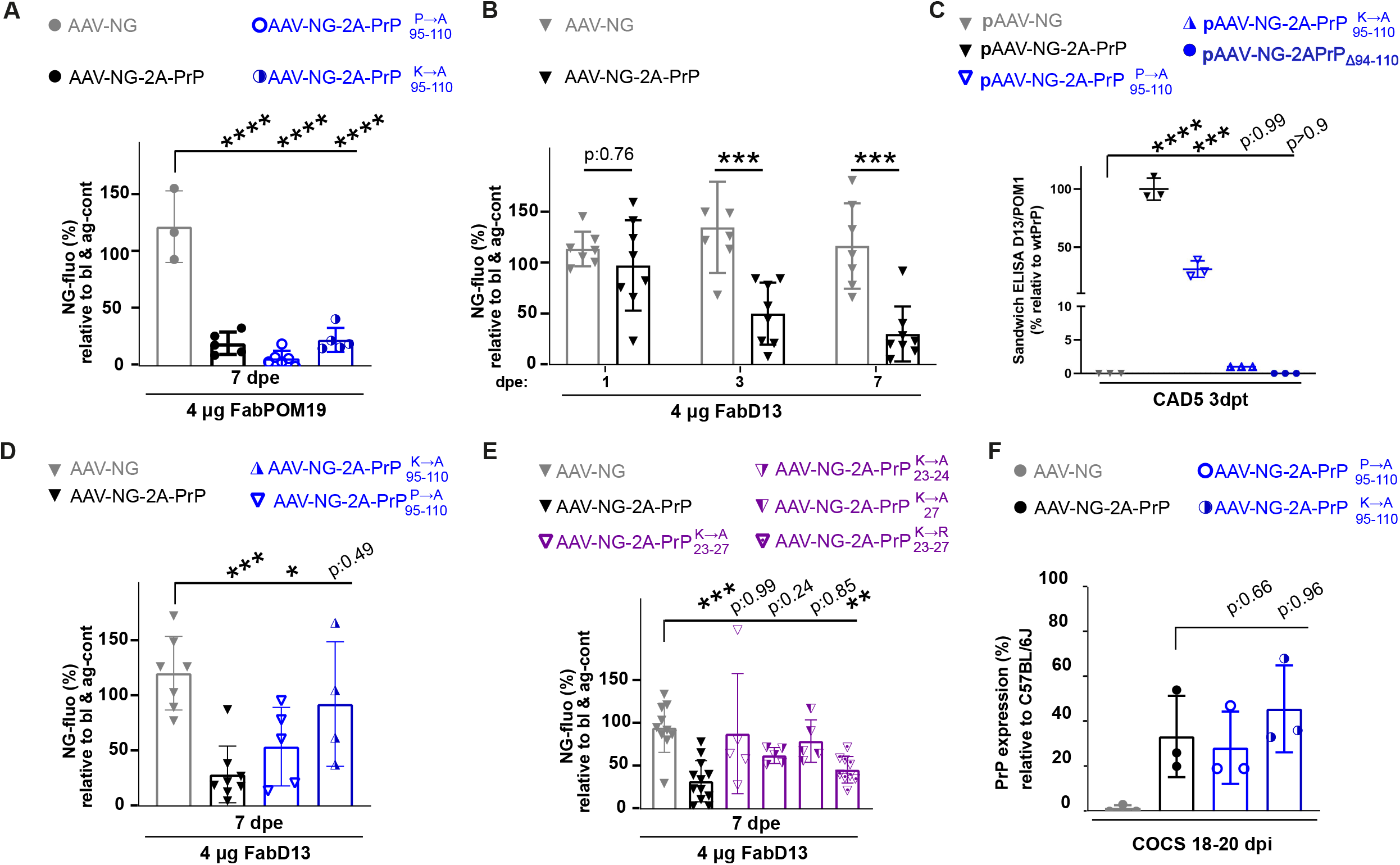
CC2-ligand mediated toxicity requires the polybasic region. **(A**) Expression of 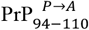 and 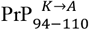 fully restore FabPOM19 mediated neurodegeneration. Here and henceforth: AAV titer 5 E+9 TU/ml, each dot represent one COCS, *P<0.05, **P<0.01, ***P<0.001, ****P<0.0001. One-Way anova, Dunnett’s post hoc test. (**B**) CC2 ligand FabD13 induced significant neurodegeneration at 72 hours and 1 week after pulse treatment of the antibody. Sidak post hoc test. (**C**) CAD5 cells were transduced with pAAV vectors and lysed after 3 days. Lysates were probed by D13/POM1 sandwich ELISA. We observed no binding of D13 to PrPΔ94-110 and drastically reduced binding to the uncharged CC2 PrP^C^ mutant 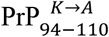. In contrast, 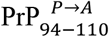 showed only minimally reduced binding to D13. All expression levels are relative to wtPrP^C^. ELISA data were performed in triplicates., **** p<0.0001. (**D**) Expression of 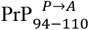 slightly sensitizes COCS against FabD13 mediated neurodegeneration in contrast to the expression of 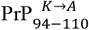. Two-Way Dunnett’s post hoc test. (**E**) *Prnp* variants with K->A mutations of the entire polybasic region, of K23 and K24, or of K27 (but not K->R mutations) fail to restore FabPOM19 toxicity. (**F**) Similar expression levels of wt and CC1-mutant PrP determined in COCS transfected together with those used for the experiments in A and D but grown separately on an inlay. Sandwich ELISA (coating: POM1; probing biotinylated POM19). Each dot represents one inlay with 11 slices.

In order to study pathological CC2 ligand interactions, we performed pulse treatment experiments with the high-affinity CC2 domain ligand D13, previously found to induce neurodegeneration with a delayed kinetic in intact mice [10]. We found significant reduction of NG expression, detectable at significant levels after 72 hours post exposure (**Fig 3B**), which represent a delay in contrast to GDL-mediated neurodegeneration (neurodegeneration 24 hours post exposure). We then investigated using ELISA the binding capacity of FabD13 to the protein mutants with an uncharged CC2 or an altered SH3 (**Table 1**). We found that the lysine residues in the CC2 largely impart FabD13 binding (**Fig 3C**). Interestingly, the SH3 mutant protein still partially mediates FabD13 induced neurodegeneration (**Fig 3D-F**).

We then asked whether the CC1 domain mediates the CC2-related toxicity in addition to GDL-induced neurodegeneration. As for GDL-mediated neurodegeneration, we found that PrP_*K*→*A* 23−27_ as well as PrP_*K*→27_ does not mediate FabD13-induced neurodegeneration (**Fig 4E**). However, in contrast to FabPOM19 induced toxicity, PrP_K23A,K24A_ was unable to fully rescue the toxicity induced by CC2 ligands (**Fig 4E**) (p = 0.24).

### PrP^C^ with an uncharged CC1 suppresses neurodegeneration induced by HC-deleted PrP^C^

Expression of PrP^C^ with a deletion of amino acid 105-125, spanning six residues of the CC2 and fifteen of the HC, is well known to induce spontaneous degeneration of the nervous system [11, 12]. Specifically, the brain of mice lacking the entire HC with an intact CC2 (Δ111-134) shows white-matter vacuolation and astrogliosis in cerebellum, brain stem and corpus callosum, without description of significant neuronal degeneration [26]. The low expression of transduced PrP_Δ111-134_ described above suggests a neurotoxic effect. To discriminate between defective expression of the viral vector and degenerative effects induced by PrP_Δ111-134_, we co-incubated slices with AAV-Syn1-NG-2A-PrP_Δ111-134_ and AAV-Syn1-NG a 1:10 ratio. In contrast to co-incubation with the AAV-Syn1-NG-2A-PrP_wt_ we found a drastic reduction of NG expression when PrP_Δ111-134_ was co-expressed with AAV-Syn1 -NG (**Fig 5A-B**), suggesting neurotoxicity of the HC truncated construct. A central characteristic of this toxic deletion mutant is that the effect can be rescued by co-expression with molar excess of wt PrP^C^ [12, 14]. We therefore co-infected COCS with AAV-Syn1-NG-2A-PrP_Δ111-134_ and AAV-Syn1-NG-2A-PrP_wt_ at a 1:10 ratio and could partly rescue the degenerative effect (**Fig 5A-B**).

**Figure 5:**
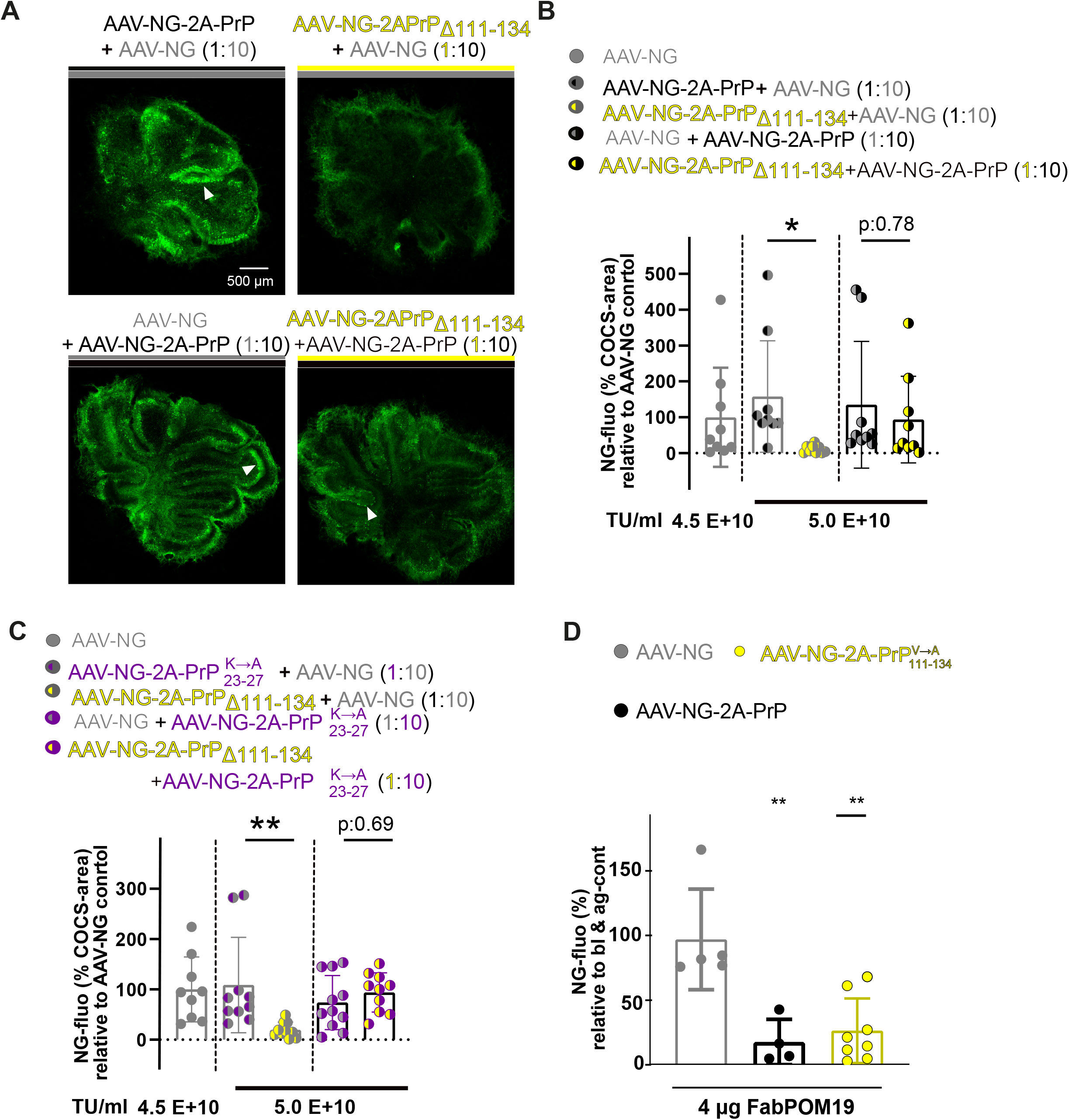
Co-expression of wild-type PrP counteracts the degeneration induced by PrP_Δ111-133_. **(A**) Representative micrographs of COCS two weeks after co-transduction with various combinations of AAV vectors. Co-transduction with 10:1 or 10:1 AAV-hSyn1-NG (4.5 E+10 TU/ml) + AAV-hSyn1-NG-2A-PrP (4.5 E+9 TU/ml) (left column) did not impair Purkinje cell viability. In contrast, co-expression of AAV-hSyn1-NG-2A-PrP_Δ111-134_ AAV-hSyn1-NG (1:10) resulted in decreased NG expression and drastic reduction of Purkinje cell labeling (upper right). This effect was partially mitigated by coexpressing AAV-hSyn1-NG-2A-PrP_Δ111-134_ + AAV-hSyn1-NG-2A-PrP (1:10; lower left). White arrowhead: NG-labeled Purkinje cell layer. (**B**) Quantification of NG expression from the experiments shown in panel A. First column NG expression after incubation 4.5 E+10 TU/ml of AAV-hSyn1-NG alone. Here and henceforth: graphs show percentage of NG^+^ pixels within COCS, divided by COCS infected with AAV-hSyn1-NG. Each dot represents one COCS, *P<0.05, **P<0.01, ***P<0.001, ****P<0.0001. Two-Way anova, Sidak post hoc test. (**C**) NG morphometry showing preservation of NG expression by co-expression of PrP_Δ111-134_ and PrP^C^ mutants with uncharged CC1 (3x K->A). (**D**) PrP mutants in which all valine residues in the HC were replaced by alanine residues restored antibody-mediated neurodegeneration. One dot represents one COCS, mean±sd, ns=not significant, 5 E+9 TU/ml; **P<0.01, one-way anova, Dunnett’s post hoc test.

Introduction of an additional deletion in the polybasic region (residues 23-31) or a combinatorial expression with an uncharged CC1 abolishes the pathological inward current of toxic 105-125 deletion mutants in cell culture [11, 30, 31]. We therefore wondered if PrP_*K*→*A* 23−27_ is incapable of rescuing PrP_Δ111-134_ induced neurodegeneration in contrast to wt PrP. However, co-expression of PrP_Δ111-134_ with PrP_*K*→*A* 23−27_ at a 1:10 fully rescued NG expression, indicating that the health of transduced neurons was restored (**Fig 5C**).

The hydrophobic valine residues within the HD are contributing to the physical properties of this domain. We therefore generated a construct in which we replaced all valines between residues 111-134 with alanines (PrP_V→A 111-134_). In COCS at 19 dpi, we found no difference in NG expression compared to AAV-Syn1-NG-2A-PrP_wt_. In fact, this construct fully mediated neurodegeneration (**Fig 5D)**.

## Discussion

### A rapid ex vivo assay to study PrP-mediated neurodegeneration by reverse genetics

PrP^C^ can mediate neuronal cell dysfunction and death in prion infections, by way of interaction with aggregated Aβ, α-Synuclein and tau, and upon exposure to antiprion antibodies [5-7, 9-11, 15]. The prevailing view is that under physiological conditions, the GD prevents pathological effects that are allosterically mediated by the FT [9, 11]. Accordingly, mice expressing an FT directly fused to the GPI anchor, but lacking the GD, develop lethal neurodegeneration [21]. Nuclear-magnetic resonance spectroscopy has revealed that the FT is entirely unstructured [1], which suggests that it may represent an optimal target for therapeutic ligands as it offers a good spacing of the amino acid residues with minimized steric effects. Data from genetically modified mice expressing PrP^C^ deletion mutants have provided important details about the functional domain of the FT [18]. We have now expanded this work with a reverse-genetics study addressing the role of various PrP^C^ domains and their composition in mediating prion-related neurodegeneration.

These studies are often performed by generating transgenic mice, which is laborious, lengthy and very costly. We have therefore developed an ex vivo method relying on AAV-mediated gene transfer of a polyprotein vector system to express PrP^C^ in combination with NG to cerebellar organotypic slices under the hSyn1 promotor. We expected transfected neurons to lose NG expression upon degeneration. In order to validate the predictive value of this method, we measured the effect of toxic antiprion antibodies on NG expression using calbindin morphometry and PI incorporation. Consistent with the experiment with transgenic PrP_Δ32-93_, which were resistant against GDL in previous experiments, we did not observe neurodegeneration at moderate PrPΔ32-93 expression levels (20% relative to the PrP concentration in wt brain homogenates), whereas deletion of the CC2 had no effect on GDL-mediated neurodegeneration [9]. We then proceeded to use this model system for assessing the effects of nineteen different *Prnp* mutants on GDL-associated neurodegeneration.

### The OR is a non-essential but powerful amplifier of allosteric FT-dependent neurotoxicity

The OR region of PrP^C^ plays multiple roles in neurodegeneration. The expansion of the OR in humans results in autosomal-dominant prion disease [32]. Transgenic mice expressing constructs with expanded OR develop ataxia and cerebellar atrophy [33, 34]. Further, the OR ligand POM2 has a protective effect on GDL-induced neurodegeneration and prolongs the life of mice expressing the neurotoxic PrP_Δ94-134_ mutant [9]. Additionally, mice expressing a truncated PrP^C^ form encompassing the OR residues (PrP_Δ32-93_) were found to be resistant to GDL-mediated neurodegeneration [9].

Here we disprove the hypothesis that the FT exerts its pathological function by integrating the OR into the plasma membrane via lysine-tryptophan residues. Further, histidine residues associated with copper binding are not essential for allosteric FT neurodegeneration. We further found that drastic overexpression of PrP_Δ94-134_ can induce GDL-mediated neurodegeneration albeit with a slower kinetic. This finding aligns with initial results from prion inoculation studies, which showed that these mice displayed milder signs of neurodegeneration compared to wt mice. Only the motor neurons in the cervical segment of the spinal cord were found to be affected [35]. Collectively, these data indicate that while the OR may not be essential, it is an important amplifier of FT-dependent neurodegeneration.

### Lysine 27: a priority target for neuroprotective compounds

In line with previous reports [11], we provide strong evidence that the polybasic region of PrP^C^ is the main effector domain of allosteric FT neurodegeneration. Mice expressing physiological levels of *Prnp*_Δ23-31_ showed no clinical signs of disease at >400 dpi after intracerebral prion (RML) inoculation [36] and mice, which express mouse PrP with mutation in the 3 lysines of the polybasic region, are highly resistant to RML and 22L prions [37, 38]. Our study provides additional evidence that the charged lysine at position 27 has a major impact on GDL-mediated neurodegeneration. It is likely that charged lysines in the polycationic cluster are the residues that mainly mediate the inappropriate interaction with membrane constituents (receptors) exerting the toxic function. Of note, the three lysines are conserved among prion proteins from at least 48 species, but are lacking in the prion protein encoded in turtle, chicken, frog and zebrafish [39]. The latter four species are not known to be susceptible to prion disease.

We demonstrate that CC1 ligand Fab83 [27] strongly interacts with all three lysines in the polybasic region including K27 and therefore represents an optimal compound for protection from neurodegeneration in prion disease. Surprisingly, Fab83 prevented prion-induced neurodegeneration in COCS less efficiently (p < 0.01) than the OR binder Fab8, Fab44, Fab71 and Fab100 (p < 0.0001) [27]. Treatment of prion-infected COCS with these OR binders resulted in reduced levels of the PK-resistant scrapie isoform of PrP^C^ (PrP^Sc^) in addition to neuroprotection. In contrast, Fab83 only affected neurodegeneration but not the level of PrP^Sc^. This finding suggests the existence of additional neurodegenerative pathways beyond allosteric FT activation. In addition, it may explain why transgenic mice with *Prnp*_Δ23-31_ expression levels greater than physiological levels still became terminally ill with a significant prolonged time course in control to wt PrP^C^ overexpressing mice (*tg*a*20*) [36]. Therefore, combining neuroprotective CC1 ligands with blockers of prion conversion might represent a rational treatment approach. Finally, blocking CC1 mediated neurodegeneration represents an interesting tool for addressing pathologic pathways independent of PrP^C^.

### A charged polybasic region is essential for CC2-mediated neurodegeneration

While antibodies and antibody-derived GD ligands can induce rapid neurodegeneration, ligands that bind to the CC2 domain can also be neurotoxic [8, 9], maybe mimicking the pathologic interaction of amyloid-β with PrP^C^. Here we show that, similarly to toxic GDL, this destructive interaction requires a charged polybasic region. However, in contrast to GDL-mediated neurodegeneration, all lysine residues of CC1 have an impact on the degenerative interaction. This may be due to differences in the structural rearrangement of the FT upon binding to CC2 ligands in contrast to GDLs. Indeed, CC2 binders have found to have the strongest stimulation on proteolytic shedding by ADAM10, associated with an extended N-terminal conformation [40]. This difference could also be the basis for the delayed kinetic of CC2 mediated neurodegeneration described in intact mice [10] and recapitulated here in COCS.

### A versatile model for prion-related neurodegeneration

Mice lacking the entire HC (Δ111-134) suffer from extensive white matter degeneration [12, 14]. Here we found that overexpression of PrP_Δ111-134_ induces degeneration of Purkinje cells, which can be rescued by co-expression of wt PrP^C^. There is evidence that a charged CC1 mediates the pathology caused by *Prnp* deletion mutants [11, 30, 31], suggesting similarities to GDL-mediated neurodegeneration. When co-expressed, wt PrP^C^ may block pathological interactions of PrP_Δ111-134_ with unknown partners through its CC1 lysines. However, also the co-expression of PrP^C^ with an uncharged CC1 prevented the neurodegeneration induced by PrP_Δ111-134_. As PrP^C^ is not required for neuronal survival, these findings suggest that the uncharged CC1 may act as an inverse agonist of the neurotoxic interaction with hitherto undefined partners, being able to engage with them but not to activate deleterious downstream cascades.

### Limitations and prospects of this study

Prion infection of cerebellar organotypic cultured slices has proved extremely useful for elucidating many aspects of the pathogenesis of these diseases. However, reverse-genetics studies have proved difficult since COCS are refractory to many types of genetic manipulations. In this study, we have found that the utilization of AAV vectors can enable a rapid analysis of the consequences of expression of many *Prnp* mutants. Crucially, AAV-mediated gene transduction has allowed us to perform experiments that in the past would have required the generation of transgenic mouse lines – an expensive and time-consuming procedure. Although COCS are ultimately derived from live mice, there use abides by the 3R precepts of animal experimentation (reduce, replace, refine) since it minimizes the number of animals necessary for drawing robust conclusions.

The AAV6 capsid infects preferentially cerebellar Purkinje neurons [41]. This peculiarity was advantageous for the current study since Purkinje neurons are large and aligned along a single plane. This morphology greatly facilitates the assessment of neurodegeneration. On the other hand, neurodegenerative diseases, including prion diseases, often display conspicuous selectivity in the type of affected cells. Hence, it is likely that the findings reported here might not be generalizable to all neuronal cell types. The availability of novel AAV serotypes and of many different promoters specific for neuronal subpopulations provides a convenient path towards modeling neurodegeneration in organotypic cultures and could be used for studying many questions of prion science and beyond.

## Contributions

**R.R.R**. designed, supervised and coordinated the research and developed the assay, performed validation experiments of the assay with the assistance of Martina Puzio (M.P) and Antonella Rosati (A.R.), cloned the polyprotein vector and the vectors for expression of mutant PrP^C^ (70%) with A.R. (30%), determined the functional AAV titer after production (60%) with A.R. (40%), checked expression of mutated prion protein in cells (30%) together with A.R. (70%), modified a previously established ELISA protocol for determining protein expression level together with Marc Emmenegger (M.E.), performed anti-prion antibody treatment and imaging of COCS cultured on cover-slips (50%) with A.R. (30%) and M.P. (20%), analyzed data and wrote the manuscript with A.A. **M.P**. performed validation experiments of the assay, performed anti-prion antibody treatment and imaging of COCS (20%) and analyzed data. **A.R**. performed validation experiments of the assay, helped with the cloning of the vectors for expression of mutant PrP^C^ (30%), determined the functional AAV titer (40%), determined protein expression level by ELISA, produced and transduced COCS, performed anti-prion antibody treatment an imaging of COCS cultured on cover-slips (30%). **M.E**. modified a previously established ELISA for determining protein expression level and edited the manuscript. **B.L.S and P.V**. produced viral vectors. **D.H. and A.C**. performed molecular dynamic experiments and analyzed data. **A.A**. conceived the primary idea of the project, advised on establishing the COCS assay, coordinated the research tasks, appropriated the funding, offered feedback and mentoring, and wrote and edited the paper.

## Acknowledgements

We thank Livia Takàcs, Ahmet Varol and Linda Irpino for technical help, Dr. Assunta Senatore for providing Fab83 and for discussions, Dr. Simone Hornemann and Prof. Ben Schuler for advice on prion protein modification, Prof. Elisabeth Rushing for editorial assistance, Rita Moos for production of the Fab fragments and Mirzet Delic for handling mice.

## Funding

A.A. is supported by institutional core funding by the University of Zurich and the University Hospital of Zurich, the Schwyzer Winiker Stiftung, and the Baugarten Stiftung (coordinated by the USZ Foundation, USZF27101), the Innovation Fund of the University Hospital Zurich (INOV00096), the Driver Grant 2017DRI17 of the Swiss Personalized Health Network (SPHN), a Distinguished Scientist Award of the NOMIS Foundation, an Advanced Grant of the European Research Council (ERC Prion2020 No. 670958) and grants from the GELU Foundation, the Swiss National Science Foundation (SNSF grant ID 179040 and grant ID 207872, Sinergia grant ID 183563), the HMZ ImmunoTarget grant, the Human Frontiers Science Program (grant ID RGP0001/2022), the Michael J. Fox Foundation (grant ID MJFF-022156), and the Innosuisse Innovation project 100.020 IP-LS.

RRR was supported for this work by a Career Development Award from the Stavros Niarchos Foundation.

## Material and Methods

### Animals

All animal experiments were conducted in strict accordance with the Swiss Animal Protection law and dispositions of the Swiss Federal Office of Food Safety and Animal Welfare (BLV). Animal protocols and experiments performed in this study were reviewed and approved by the animal welfare committee of the Canton of Zurich: ZH90/2013; ZH139_16. We used the C57BL/6J inbred strain and mice from the following genotypes: Zurich III *Prnp*°/° (*Prnp*ZH3/ZH3) [19] and *tg*a*20* [42].

### Cell lines and chemicals

CAD5 is a subclone of the central nervous system catecholaminergic cell line CAD [43]. Generation of the CAD5 *Prnp*^-/-^ was previously described [44]. HEK 293T was derived from ATCC. Cells were cultured in DMEM supplemented with 10 % FBS, 1% GM and 1% PS at 37 °C in a humidified atmosphere with 5% CO_2_. All compounds were purchased from Sigma/Aldrich unless otherwise stated.

### Polyprotein vectors and adeno-associated virus (AAV) production

A polyprotein vector with a P2A sequence (GSGATNFSLLKQAGDVEENPGP) was assembled using Golden Gate cloning [45]. Using the Esp3I (Thermo Scientific) enzyme, the sequence of the NG (mNG) fluorophore, P2A sequence and murine wt PrP^C^ were assembled into the pCAG-T7 Golden Gate assembly destination vector (pCAG-TALEN-Destination, a generous gift from Dr. Pawel Pelczar). The NG-P2A-PrP^C^ cassette was then subcloned into adeno-associated virus (ssAAV) vector backbones with AAV2 inverted terminal repeats (ITRs), kindly provided by Bernhard Schneider (EPFL, Switzerland). In this vector, the expression of the cassette is driven by the human Synapsin I (hSyn1) promoter. All further vectors for the expression of mutant PrP^C^ were produced by cloning a synthetic gene block (gBlock, IDT, full sequence was deposited on FigShare, https://doi.org/10.6084/m9.figshare.c.5893787.v3) between the BsrGI and HindIII site of the vector replacing the wt PrP^C^ sequence.

Viral vectors and viral vector plasmids were produced as hybrid AAV2/6 (AAV6 capsid with AAV2 ITRs), contributed by Bernhard Schneider (EPFL, Switzerland) and the Viral Vector Facility (VVF) of the Neuroscience Center Zurich (Zentrum für Neurowissenschaften Zürich, ZNZ, Switzerland). The identity of the packaged genomes was confirmed by Sanger DNA-sequencing (identity check).

### Determination of the functional AAV titer

The functional (infectious) titer in transduction units (TU/ml) was determined by qPCR after S1 nuclear digestion [46]. Viral titration was performed in HEK 293T cells. As a reference, additional HEK 293T cells were infected with an AAV-cmv-eGFP-WRPE virus with a known titer established by flow cytometry (kindly provided by Bernhard Schneider, EPFL). DNA was extracted from the cell homogenate using a NucleoSpin tissue mini kit (Macher-Nagel; ref 740952.5) following the manufacturer’s instructions. After S1 nuclear digestion (promega, M5761), qPCR was performed for the woodchuck hepatitis virus post-transcriptional regulatory element (WRPE, forward primer: CCG TTG TCA GGC AAC GTG; reverse primer: AGC TGA CAG GTG GTG GCA AT) and albumin (forward primer: TGA AAC ATA CGT TCC CAA AGA GTT T; reverse primer: CTC TCC TTC TCA GAA AGT GTG CAT AT) including standards with a known concentration for WPRE and human albumin. Based on these standards, the viral DNA copy number per cell was established by dividing copy number WRPE over the copy number for albumin. Finally, TU/ml was established by linear regression and correlated with the reference virus.

### Antibodies and recombinant PrP generation

POMs [47] and D13 [48] mouse monoclonal antibodies were produced using the hybridoma technology. Purification was performed by affinity chromatography using protein G sepharose, diluted in PBS. Antibody purity was assessed with silver-stained SDS-PAGE gels. Recombinant mouse PrP derived from E. Coli was purified by affinity chromatography, oxidized on-column and then refolded into the native state [49]. F(ab) POM fragments were generated in-house from the POM antibodies using papain digestion and purified with Protein A agarose, followed by size exclusion chromatography.

For staining and Western blotting, we purchased the following primary antibodies: Anti Calbindin rabbit (Abcam Ab25085, immunostaining 1:500); Anti Myelin oligodendrocyte glycoprotein antibody (Abcam, MOG, Ab32760, immunostaining 1:200); Anti Neuronal Nuclear antibody (Millipore, NeuN, A60, immunostaining 1:500 and Cellsignal, D3S3I, Western blotting 1:500); Anti glial fibrillary acid protein (Abcam, GFAP, Dako Z0334 1:1000); Anti CD68 (Biorad MCA1957, immunostaining 1:500); Anti GR778/BIP (Abcam, immunostaining 1:500); anti-α-GAPDH (Sigma-Aldrich, 1:15’000). The following secondary antibodies: Alexa 546 goat anti-mouse; Alexa 647 goat anti-rabbit, Alexa 594 goat anti-rat (all Biolegend, 1:1000), and horseradish peroxidase (HRP)-goat anti–rabbit IgG1 (1:10,000, Zymed) were used for staining and western blotting.

### Lipofection of cells

CAD5 *Prnp*^-/-^ cells were cultured in a 12-well format on gelatin coated (2% Gelatin) coverslips or in 6-well plates until they reached approximately 90% confluence. Cells were then transfected with the plasmid using Lipofectamine® 2000 (Invitrogen) according to manufacturer’s instructions.

### Immunohistochemical staining of cells

Cells grown on cover slips were fixed in 4% paraformaldehyde, then incubated for 10 minutes in TritonX solution (0.1% in 0.5% bovine serum albumin dissolved in PBS) and blocked with 0.5% bovine serum albumin for 45 minutes. The cells were incubated with primary antibody (POM19: 2 μg ml-1; BIP 1:500) and diluted in PBS containing 0.5% bovine serum albumin for 1 hour. After washing, the cells were incubated with a fluorescent-labeled secondary antibody (Alexa Fluor 546 anti-mouse and Alexa Fluor 647 anti-rabbit; 1 μg ml-1). In the last PBS wash, 4,6-diamidino-2-phenylindole, dihydrochloride (DAPI; Molecular Probes) was added and sections were mounted with Fluor Save (Calbiochem). For analysis, images were acquired with a FluoView® FV10i Confocal Laser Scanning System.

### Production, AAV transduction and culturing of cerebellar organotypic slice cultures (COCS)

Cultured Organotypic Cerebellar Slices (COCS) from *Prnp*^ZH3/ZH3^ and *tg*a*20* mice were obtained as previously published [50]. 350 μm thick COCS were prepared from 9-12 day-old pups. After preparation, COCS were infected with AAV on a free-floating section directly after production on day 0. 21. COCS were incubated in a 6-well plate with AAV at a final concentration between 1 E+9 and 1 E+10 (as specified in the figure legend) TU/ml diluted in physiological Grey’s balanced salt solution for 1h at 4°C on a shaker.

From every incubation, 11 COCS were placed and cultured on a Millicel-CM Biopore PTFE membrane insert (6-well inserts, Millipore) in order to determine the PrP^C^ expression level based on a sandwich enzyme linked immunoabsorbent assay (ELISA; see below). The remaining 10 COCS were placed on cover slips and cultured in roller tubes following the published protocol [20]. In brief, coverslips were surface coated with poly-D-lysine (30’000-70’000 mol, Sigma) solution overnight. Following the addition of 30-μl of 100 U/ml thrombin (Merck) in Grey’s BSS to coagulate the plasma by stirring with a pipette tip, COCS were then placed in a 20-μl drop of chicken plasma (Cocalico). The clot was allowed to dry for 3-5 minutes. Coverslips were then placed in a screw-top, flat-sided culture tube containing 750 μl of culture media. Subsequently, the tubes were placed in a roller drum-housed incubator at 36°C with 10 revolutions/hr to allow proper aeration and feeding. The roller drum was tilted at 5° to ensure that slices received the medium during half of each rotation.

### Anti-prion antibody treatment, imaging and quantification of COCS cultured on cover slips

After 19±5 days in culture, COCS cultured on cover slips were scanned using the Olympus FLUOVIEW FV1000 confocal laser scanning microscope. Three slices were scanned at the same time. Coverslips were removed from the roller tube using a Fine Point High Precision Forceps (Fisherbrand™) and then placed in a petri dish with a glass bottom and support with a drop of PBS. Slices were scanned at the following properties: Z moving range: -100 - 2’800, enabling overlap stage position, upper (−400,500) AR searching range, high speed (256×256) map image resolution for each area, 0.17 mm of correcting ring at 38°C. Using the Z-stack multi area, 5 images per slices were taken at +50, 25, -50 (see also Supplement Figure 4). The confocal aperture was set at x5 and the laser was set at 10% with a fix sensitivity (regularly at 43%) over the whole experiment, in dependence of the mNG fluorescence expression level. From the 5 stack images, maximal intensity projections were generated using Fiji and images were converted into grey-scale images. Using the image analysis software “cell^P” (Olympus) the region of interest (ROI) and the COCS area were drawn in the overlay. Within this ROI percentage of NG expression (percentage of pixel) was determined with identical greyscale threshold setting for identifying positive pixels.

After baseline imaging, 4 μg of Fab fragments of anti-prion antibodies or 4 μg of Fab fragment of anti-prion antibodies pre-incubated with a three-molar excess of recombinant PrP^C^ in 10 μl of PBS were directly dropped on the slices. Imaging with Fluoview was repeated (24 hours, 72 hours) 1-week after exposure to the antibody. The expression level was then calculated in relation to baseline expression.

### Generation of Homogenate from CAD5 culture or COCS for protein analysis

To produce the homogenate, 100 μl of cOmpleteTM Mini EDTA-free Protease Inhibitor Cocktail (Sigma Aldrich) were added to 800 μl of Lysis Buffer pH 7.5 (RIPA buffer). CAD5 cells grown in a 6-well plate were washed with PBS, scraped from the plate and aspirated in PBS solution. After centrifugation. homogenization was performed by adding lysis buffer as well as physical lysis with an 18-gauge syringe. COCS gown on PTFE Millicells inserts were washed twice in PBS and scraped off the PTFE membranes with PBS. Homogenization was performed with a TissueLyser LT (Qiagen) for 2 minutes at 50 Hz in lysis buffer. Bicinchoninic acid assay (Pierce™ BCA protein assay kit, Thermo Fisher Scientific) was used to determine protein concentrations.

### Protein analysis (ELISA and Western blotting)

The enzyme linked immunoabsorbent assay (ELISA) was modified from a previously established protocol [47, 51] to a sandwich ELISA protocol for PrP^C^ detection in 384 well plates. All washing, aspirating and dispensing steps were performed with the BioTek Washer EL406 and MultiFlo FX. Plates were coated at 4°C overnight with 0.4 ng/μl POM1 (or 7.5 ng/μl Fab83 in SFig.11H or 0.4 ng/μl D13 in SFig12H) in sodium carbonate and were washed 5 times with PBS containing 0.1% (vol/vol) Tween-20 (PBS-T) the next morning. Subsequently, they were blocked with 5% SureBlock™ (LuBioScience GmbH) in PBS-T and then incubated for two hours at 25 °C. In the next step, the blocking solution was aspirated and the plates were incubated again for two hours at 25°C with 20 μl of of either rmPrP23-231 or 2-fold serially diluted slices homogenate (prediluted to a concentration of 2000 μg/ml total protein) in PBS-T containing 1% SureBlock™. Plates were washed 5 times with PBS-T and incubated for one hour at 25°C with 20 μl of biotinylated POM19 (used at 1:10000 dilution in 1% SureBlockTM, PBS-T). After washing the plates again with PBS-T, they were probed with horseradish peroxidase-Streptavidin (BD Biosciences; used at 1:1000 dilution in 1% SureBlockTM, PBS-T). Following the final washing step, plates were incubated for 10 minutes with TMB stabilizing chromogen (Invtirogen) and developed using 0.5 M sulphuric acid. The optical density was measured at 450 nm using an EnVision Platereader. Using the rmPrP standard curve, we interpolated the values of the brain homogenates to calculate PrP concentration per sample. Following these steps, the PrP concentrations obtained for each sample were normalized to the respective concentrations of the tissue homogenates of Bl6 (positive control) and ZH3 (negative control) mice in order to counterbalance potential assay-to-assay heterogeneity. We thus show PrP levels as percentages relative to Bl6 PrP expression.

For analysing lysate under deglycosylation conditions, homogenates were incubated with PNGaseF (NewEngland Biolabs) following manufacturer’s instructions. For Western blotting, equal amounts (25 μg protein) of cell or COCS homogenate with or without PNGaseF digestion were mixed with loading dye (ThermoScientific) and loaded on a 4-12% precast NuPage gel (Invitrogen). Transfer to a PVDF membrane (Invitrogen) was performed using a iBlot™ Cel Transfer Device. PrP^C^ was detected by Western blot using the monoclonal antibody POM19 at 0.1 μg mL^-1^. NeuN was detected by using anti-Neun (D3S3I) antibody. Loading control was performed with anti-α-GAPDH (Sigma-Aldrich, 1:15’000). Horseradish peroxidase (HRP)-goat anti–rabbit or mouse IgG1 (1:10,000, Zymed) was used as secondary antibody. Membranes were illuminated using a Stella Imaging System.

### Immunostaining of COCS

COCS grown on PTFE were washed twice in PBS and fixed in 4% paraformaldehyde for at least 2 days at 4°C and were washed again twice in PBS prior to blocking of unspecific binding by incubation in blocking buffer (0.05% Triton X-100 vol/vol, 0.3% goat serum vol/vol in PBS) for 1 h at room temperature. The primary antibody was dissolved in blocking buffer and incubated for 3 days at 4°C. After three washes with PBS for 30 min, COCS were incubated for 3 days at 4°C with fluochrome conjugated secondary antibodies at a dilution of 1:1’000 in blocking buffer. Slices were then washed with PBS for 15 min and incubated in DAPI (1 μg mL^-1^) in PBS at room temperature for 30 min to visualize cell nuclei. Two subsequent washes in PBS were performed and COCS were mounted with fluorescence mounting medium (DAKO) on glass slides. Pearson’s correlation coefficient was calculated using the macro JACoP [52] after background subtraction in Fiji.

### Propidium iodide (PI) staining of COCS

For propidium iodide (PI, Sigma) staining of COCS cultured on cover slips, PI were diluted in slice culture media at a final concentration of 10 uM. The media in roller tubes were replaced by media containing 10 μm of PI solution and slices were incubated for 30 minutes. After incubation, the media were replaced with fresh media and returned to the incubator for 10 minutes prior to imaging with Olympus FLUOVIEW as described above.

For COCS cultured on PTFE membranes 1 ml of media containing PI solution (10 uM). PTFE membranes with COCS were placed into the solution and incubated at room temperature for 30 minutes, covered with aluminum. Subsequently, petri dishes containing stained COCS on membrane were imaged with Olympus Fluoview (excitation maximum: 534 nm; emission maximum: 617).

### Simulation details

All molecular dynamics simulations were performed using the GROMACS simulation package [53] and the CHARMM36 force field [54] with the TIP3P water model. Most simulations were carried out with the 3-octapeptide repeat (residues 60-83) of the flexible tail (FT) of PrP^C^ except for a run with residues 23-90. The DPPC (dipalmitoylphoshpatidylcholine) bilayer was prepared using the GROMACS simulation package [53]. Equilibration consisted of 10 ns. During both equilibration and production runs, van der Waals interactions were switched off at a distance of 1.0 nm and electrostatic short-range interactions were cut-off beyond a distance of 1.2 nm. The long-range electrostatic interactions were treated with Particle Mesh Ewald [55]. The simulation temperature of 323 K, which is above the main phase transition of the DPPC bilayer, was kept constant using the v-rescale algorithm [55]. The Berendsen pressure coupling algorithm [56] was employed for constant pressure simulation at 1 atm. Periodic boundary conditions were applied. All simulations started with the peptide at a distance of at least 1.5 nm from the bilayer surface. A total of four independent simulations with different initial velocities were carried out for the wt and three simulations for the triple-point mutant. The length of each run was between 170 and 410 ns (see Fig S7) and snapshots were saved every 50 ps. A time step of 2 fs was employed for all runs.

### Statistical analysis

The two-way Anova Sidak post hoc test for multicolumn comparisons or the Dunnett’s post-hoc test for comparisons of all columns to a control column were used for statistical analysis of experiments involving the comparison of three or more samples. A paired Student’s t-test was used for comparing two samples. Results are displayed as the average of replicates ± s.d. Significance is defined as a p value below 0.05.

## Supplemental Figure Legends

***Supplemental Figure 1: Expression of polyprotein constructs***

(**A**) Transduction with AAVs encoding NG (upper row) or NG-2A-PrP (lower row) (Figure 1B) in differentiated CAD5 cells. Surface immunostaining (without permeabilization) for PrP (antibody POM19) six days post lipofection. Nuclear staining: DAPI.

(**B**) Expression of the NG-2A-PrP polyprotein in cerebellar organotypic slices (COCS) 19 days post transduction with AAV-hSyn1-NG-2A-PrP. Immunostaining with POM19 (magenta). Nuclear staining: DAPI. White star: Purkinje cells. Granule cells (GC), Purkinje cell (PC), molecular layer (ML). Boxes: region magnified in Figure 1C. (**C**) Representative images of co-immunostaining for calbindin (Purkinje cells) and NeuN (cerebellar granule cells) 19 days post transduction with AAV-hSyn1-NG-2A-PrP. Nuclear staining: DAPI. Cyan: DAPI, green: NG, blue: calbindin, magenta: NeuN.

(**D**) Immunostaining of AAV-hSyn1-NG-2A-PrP-transduced COCS for CD68 (microglia). Cyan: DAPI, red: CD68. (**E**) Micrographs of immunostainings for glial fibrillary acidic protein (GFAP), expressed in astrocytes, and myelin oligodendrocyte glycoprotein (MOG), expressed in oligodendrocytes. 19 days post transduction with AAV-hSyn1-NG-2A-PrP. Nuclear staining: DAPI. Cyan: DAPI, green: NG, blue: MOG, magenta: GFAP.

***Supplemental Figure 2: Reconstitution of FabPOM19 mediated neurodegeneration in PrnpZH3/ZH3-COCS***

(**A**) Representative micrographs of fixed and immunostained COCS after treatment with FabPOM19 (10d, 134 nM, multiple doses). B We observe broad NG expression in Purkinje cells of COCS transduced with AAV-hSyn1-NG (white star, upper line), but complete loss of NG expression in COCS transduced with AAV-hSyn1-NG -2A-PrP (lower line). Representative calbindin staining, a marker of Purkinje cells, demonstrate a reduction of Purkinje cells axons (yellow asterisk) and a slightly reduced DAPI signal in FabPOM19 treated COCS transfected with AAV-hSyn1-NG -2A-PrP in contrast to AAV-hSyn1-NG. (**B**) Morphometric quantification of NG shows a significant difference between FabPOM19 treated AAV-hSyn1-NG and AAV-hSyn1-NG-2A-PrP-transduced slices. Each dot represents an individual COCS. Here and henceforth: mean ± sd, student t-test, **<0.001, ns: not significant. Data points are divided by the mean of AAV-hSyn1-NG control. (**C**) Morphometry of the same slices as analyzed in B from AAV-hSyn1-NG -2A-PrP indicates just 80% of DAPI signal in contrast to AAV-hSyn1-NG -2A-PrP transduced slices. However, the effect is not significant (p:0.13).. (**D**) Quantification of calbindin expression also shows a 20% reduction of signal but no significant (p:0.34) change between FabPOM19-treated AAV-hSyn1-NG and AAV-hSyn1-NG -2A-PrP transduced slices.

***Supplemental Figure 3: Pulse treatment of COCS***

Tga20^+/+^ COCS were cultured on inlays and treated with a pulse (4 μg/slice) of FabPOM19 optionally preincubated with a three-fold molar excess of rmPrP for 4, 8 (**A, B**), 24 and 48 hours (**C, D**). 10 μg of protein prepared from a homogenate of the COCS was then immunoblotted and probed with anti-NeuN antibody. (**B**) Statistical analysis indicates no significant decrease in NeuN expression upon FabPOM19 pulse treatment at early time points. N = 3 inlays, mean±sd, two-way Anova, Bonferroni multiple comparisons, NS: not significant. (**D**) Significant decrease in NeuN densitometry 24 and 48 hours post pulse treatment. N = 3 inlays, mean±sd, two-way Anova, Bonferroni multiple comparisons, **P<0.01, *P<0.05. (**E**) Propidium iodide (PI) staining of *tg*a*20*^+/+^ COCS cultivated on cover slips pulse treated with either FabIgG, FabPOM19 preincubated with rmPrP or FabPOM19. (**F**) Quantitative analysis of PI-stained COCS 24 hours post pulse treatment. Significantly increased staining intensity in FabPOM19-treated slices. N = 9 COCS, mean±sd, one-way Anova, Tukey multiple comparisons. (**G**) Time-course analysis of PI densitometry 24, 48 and 72 hours post pulse treatment revealing an early increase in intensity at 24 hours, which then decreases at 48 and 72 hours. N = 2-3 COCS, mean±sd, one-way anova, Dunnett’s multiple comparisons.

***Supplemental Figure 4: In vivo imaging of COCS on cover slips and intensity projections***

(**A**) Images of COCS are taken at five different depths every 25 μm. Confocal images are taken with a confocal aperture of 2.5, resulting in a slice thickness of 25 μm. The blue arrow represents the maximal intensity projection of all pictures into the ground plot. (**B**) Representative images of slices at different positions along the vertical axis. The middle point (0) is taken at the point of maximal NG brightness. (**C**) Maximal intensity projection of images shown in B. (**D**) Conversion of the maximal NG projection to grey scale and threshold on intensity. (**E**) Threshold on intensity from the AAV-hSyn1-NG -2A-PrP transduced slices shown in Figure 1E at baseline and 1, 3 and 7 days post exposure to 4μg FabPOM19. White dashed line: region of interest, which equals the COCS area. For each image NG-fluorescence in percentage of COCS area is given. (**F**) Representative NG-fluorescence two and four weeks post transduction with AAV-hSyn1-NG -2A-PrP. (**G**) NG morphometry in percentage of COCS area from 13 COCS two and four weeks post transduction with AAV-hSyn1-NG -2A-PrP.

***Supplemental Figure 5: FabPOM1 and FabPOM19 exert similar effects on COCS***

*Prnp*^ZH3/ZH3^ COCS were transduced with NG-2A-PrP cultured on cover slips (roller tubes) for 19 days and then treated with 4 μg/slice (pulse treatment) of FabPOM19 or 4 μg/slice FabPOM19 preincubated with a three-fold molar excess of rmPrP. (**A**) Representative propidium iodide (PI) staining 24 hours post treatment. Some Purkinje cells show co-localization with positive PI staining (asterisks). (**B**) PI densitometry showed a non-significant (p: 0.11) increase in signal for the FabPOM19 treated group in contrast to blocked control. N = 3 COCS, mean±sd, student t-test, ns: non-significant. (**C**) NG morphometry of COCS transduced with AAV-Syn-NG-2A-PrP or AAV-NG. Time course analysis after pulse treatment with FabPOM1 (24, 72 hours or 1 week post treatment) shows significant change between the groups at all-time points. N = 4-6 mean±sd, **P<0.01, ****P<0.0001 two-way anova, Sidak post hoc test. The data are normalized against rmPrP-blocked control.

***Supplemental Figure 6: All prion protein mutants undergo correct biogenesis and surface expression***

Representative PrP (POM19) surface immunofluorescent stainings of not permeabilized CAD5-*Prnp*^*-/-*^ cells transduced with pAAV-Syn-NG-2A-PrP mutants. POM19 immunostaining (middle panel in magenta) demonstrate surface expression of the prion protein on the soma (yellow arrowhead) and on the process (yellow star) of the neurons. NG (in green) expressed by the construct demonstrates the cytoplasm of transduced cells. All constructs show cell surface expression. Nuclear staining: DAPI (Cyan).. (**A**) Deletion mutants. (**B**) Alterations affecting the polybasic region. (**C**) Octapeptide repeat mutants, (**D**) Alterations affecting the charge cluster region or the hydrophobic core.

***Supplemental Figure 7: Molecular dynamics study***

(**A**) Summary of simulations: Tested peptides, structure of bilayer (Dipalmitoylphoshpatidylcholine: DPPC) and MD simulation system representing PrP^C^ (cyan) and the bilayer. (**B**) The PrP60-83 (part of the OR) inserts spontaneously into the membrane within a 100 ns time scale. Simulation in duplicates. (**C**) Insertion of the PrP23-90 (encompassing CC1, pre-Or and OR) within 50 ns. (**D**) Molecular dynamic study of three-point tryptophan to alanine mutant (lower row), of the PrP60-83 peptide showing no insertion of the FT into the plasma membrane in contrast to wt PrP (upper row). Simulation in duplicates.

***Supplemental Figure 8: Intact glycosylation of all prion protein mutants***

POM19PrP-Western blot before (left) and after deglyPNGase F; (right) of all PrP mutants. The wt PrP^C^ has two glycosylation sides at position of the residues 181 and 197 resulting in three possible glycosylation states. Two major bands at around 33-35 kDa correspond to the di- (three violet stars) and around 30kDa to the mono-glycosylated (two violet stars) isoform of PrP^C^. The non-glycosylated form demonstrates a band at 27 kDa (one violet star).(**A**) Deletion mutants. PrPΔ111-134 demonstrate a low level of expression in contrast to the other deletion mutants. (**B**) Alterations affecting the polybasic region. (**C**) Octapeptide repeat mutants, (**D**) Alterations affecting the charge cluster region or the hydrophobic core.

***Supplemental Figure 9: Expression level of prion protein mutants in transfected COCS***

(**A-B**) POM19PrP expression of wt and mutant PrP measured by sandwich ELISA of homogenate from COCS grown on inlay (11 COCS/inlay). The COCS used for ELISA experiments were transfected together with those used for the neurodegeneration assays but grown separately on an inlay. Transduction with AAV (titer: 10^9^ TU/ml). N = 3-4 inlays, mean±sd, ns: not significant, one-way Anova, Dunnett’s post hoc test. PrP concentrations (% relative to PrP concentration in cerebellar homogenate of 11 day old wt mice). (A) Octapeptide mutants. (B) Deletion mutants.

## Notes

https://doi.org/10.6084/m9.figshare.c.5893787.v3

### Competing Interest Statement

The authors have declared no competing interest.

