## Supplement for "Rapid ex vivo reverse genetics identifies the essential determinants of prion protein toxicity"

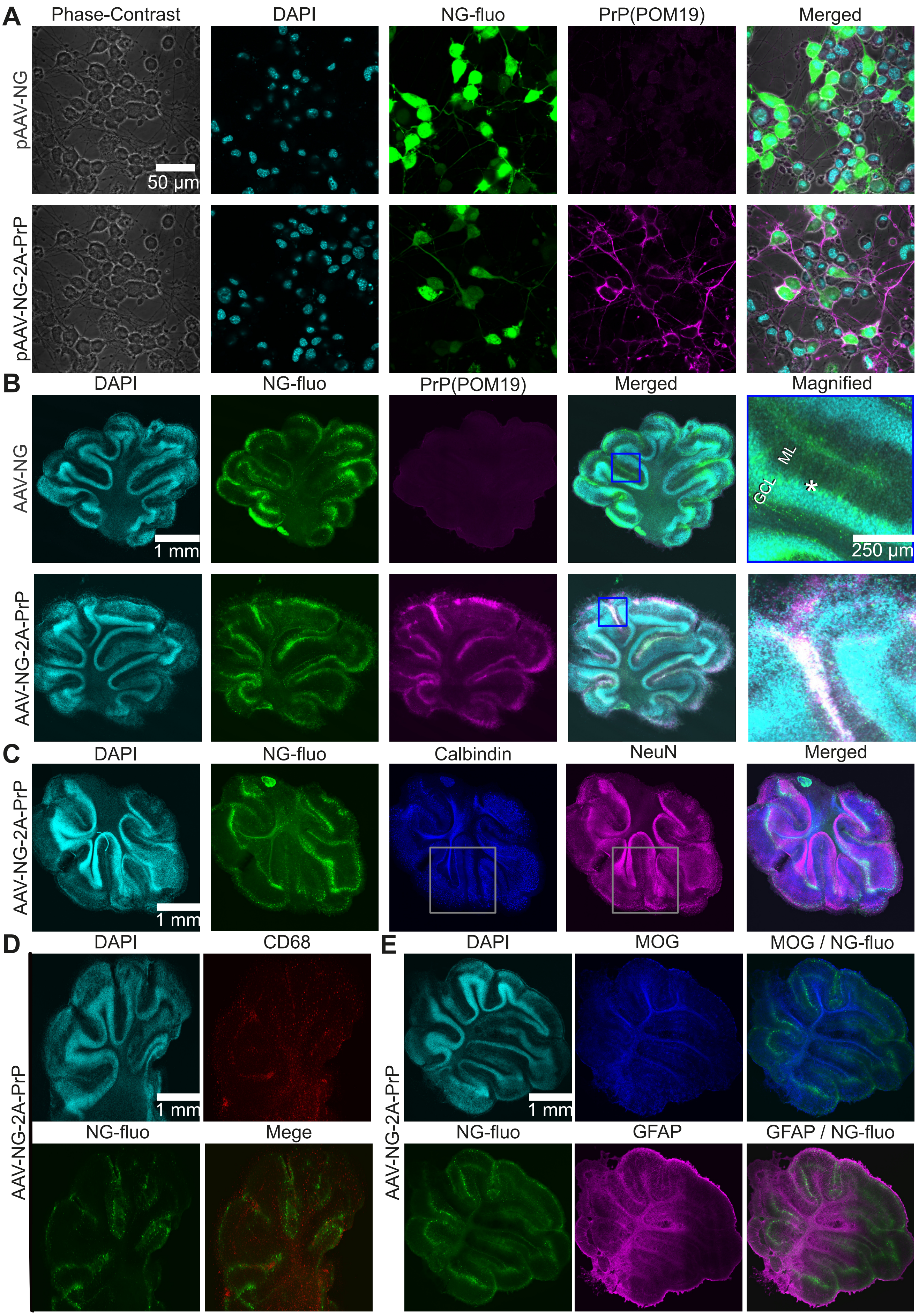


Reimann et al 2022, Supplement Figure 1

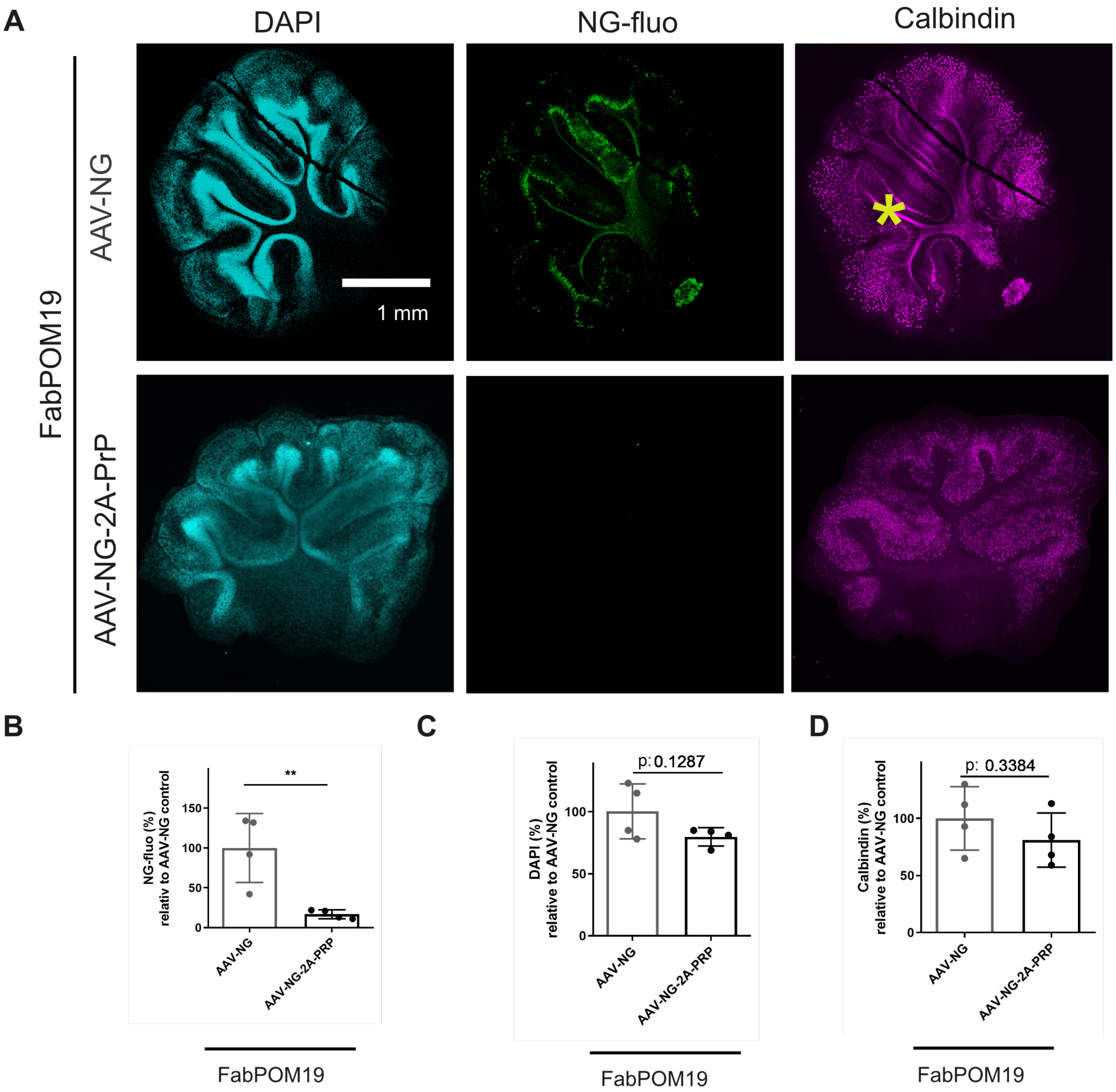


Reimann et al 2022, Supplement Figure 2

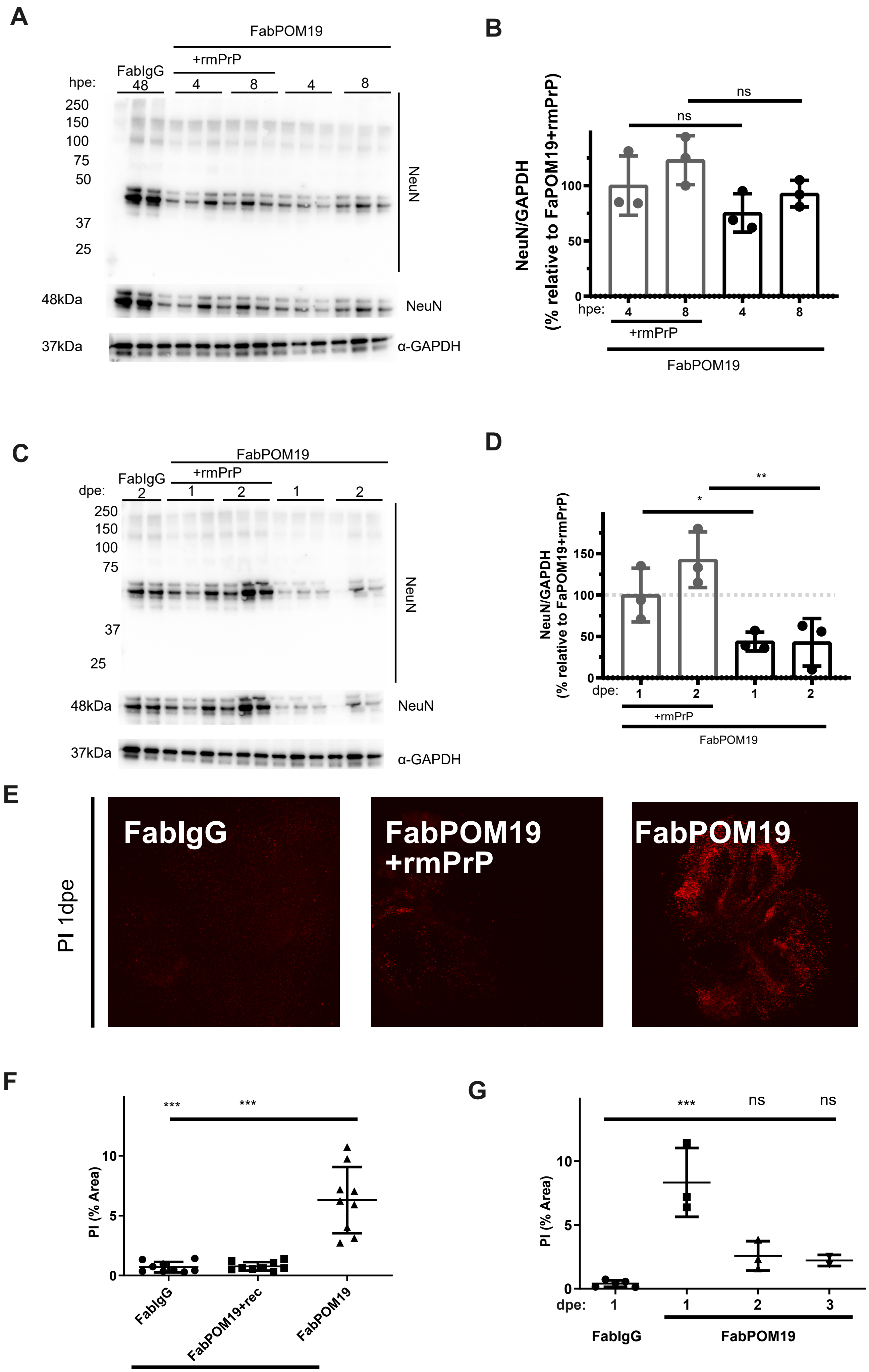


Reimann et al 2022, Supplement Figure 3

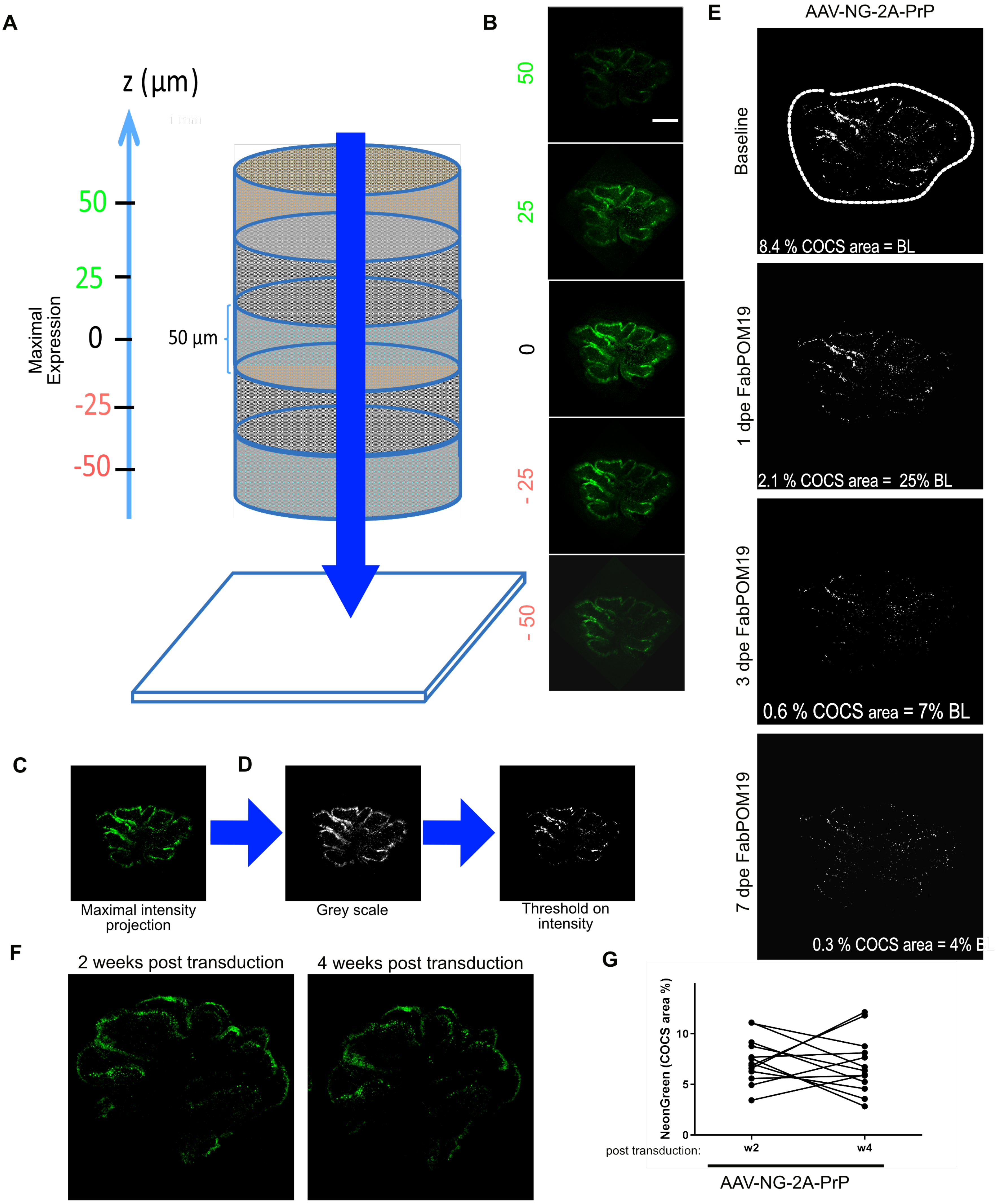


Reimann et al 2022, Supplement Figure 4

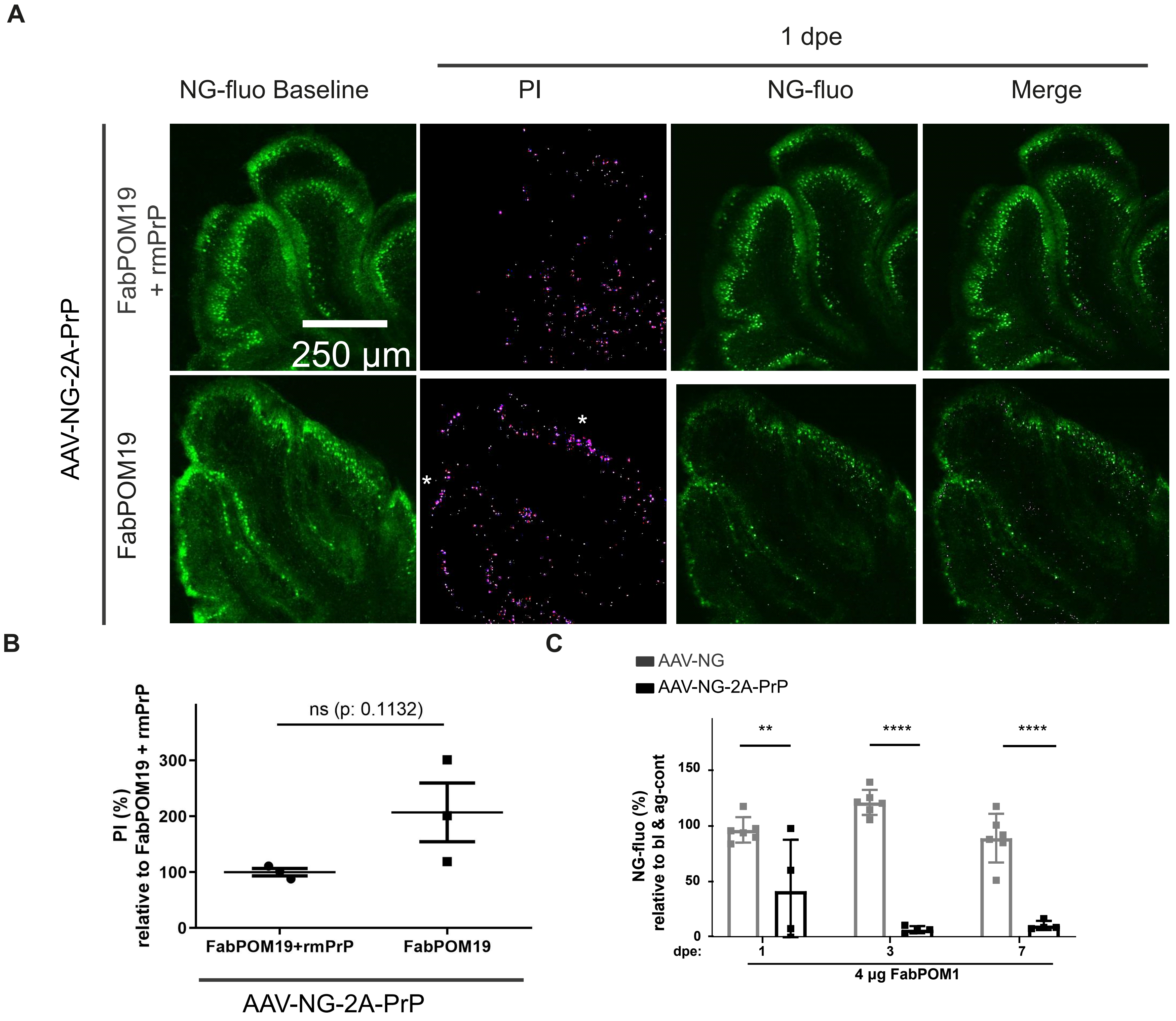


Reimann et al 2022, Supplement Figure 5

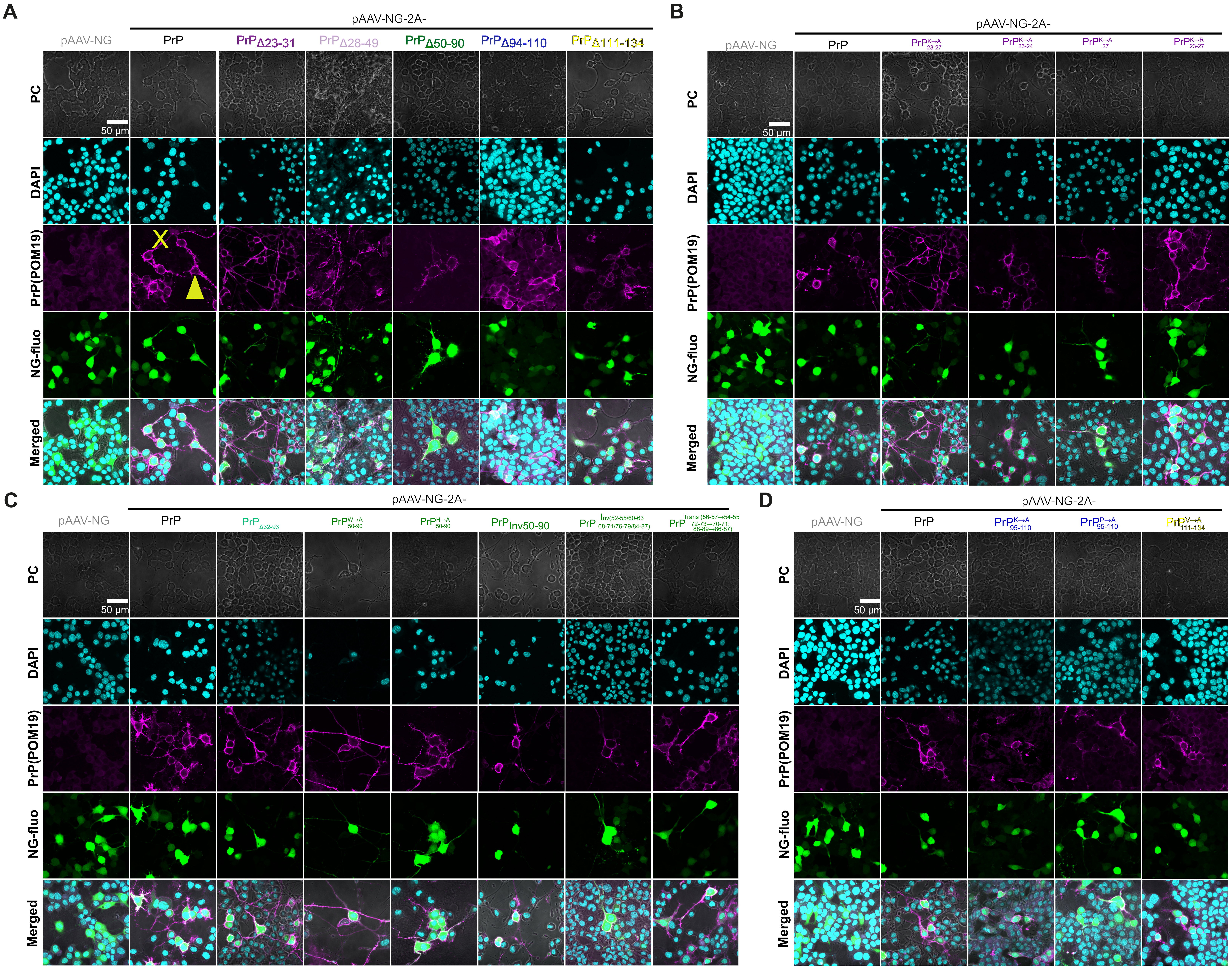


Reimann et al 2022, Supplement Figure 6

A

| Peptides | Simulation Time (ns) | lateral tension (mN/m) | Number of DPPC per layer |
|---|---|---|---|
| 1. prion 60-83 (3-octapeptide) | 389/410 | 0 | 128 |
| 2. prion 23-90 | 250 | 0 | 128 |
| 3. prion 60-83 | 390 | 0 | 64 |
| 4. prion 60-83 with Trp->Ala mutations | 170/170 | 0 | 64 |

prion 60-83 (3-octapeptide):

HGGGWGQP-HGGSWGQP-HGGSWGQP

Structure and nomenclature for DPPC

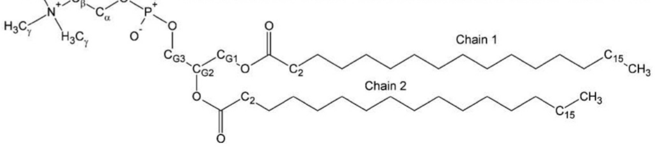


JPCB 2008, 5924

MD simulation system

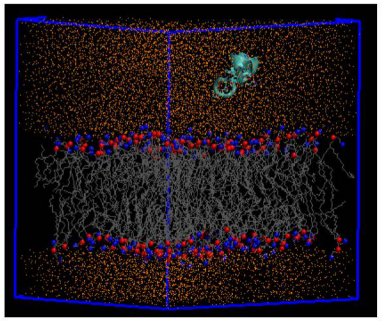

B

PrP 60-83

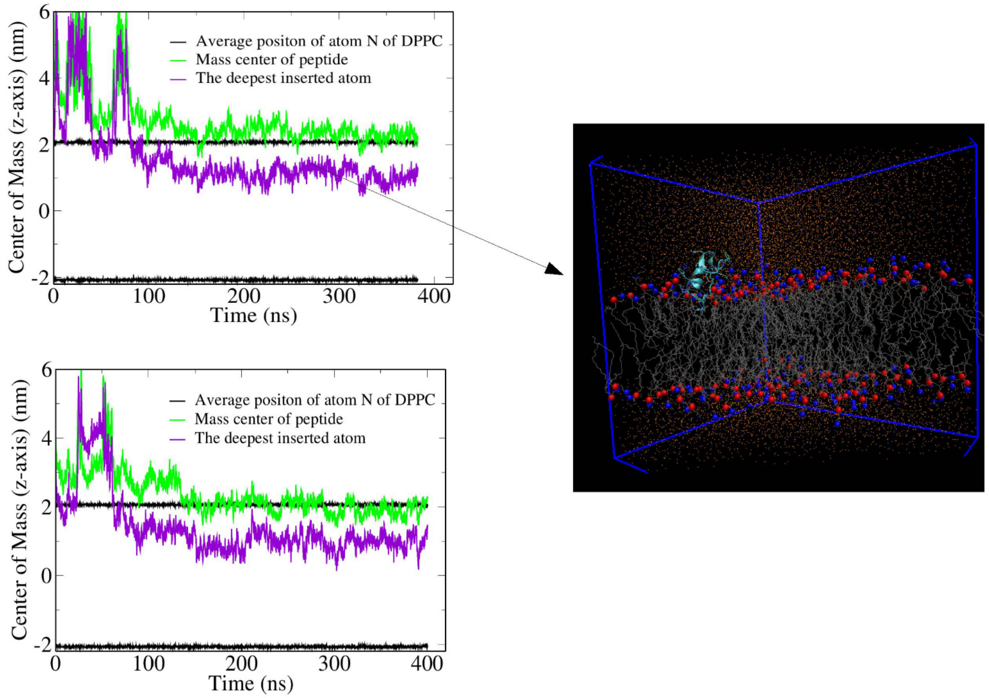


C

PrP 23-90

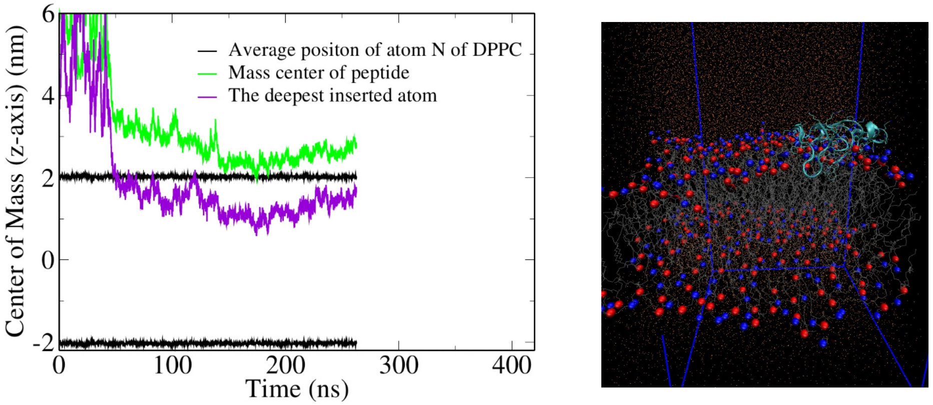


D

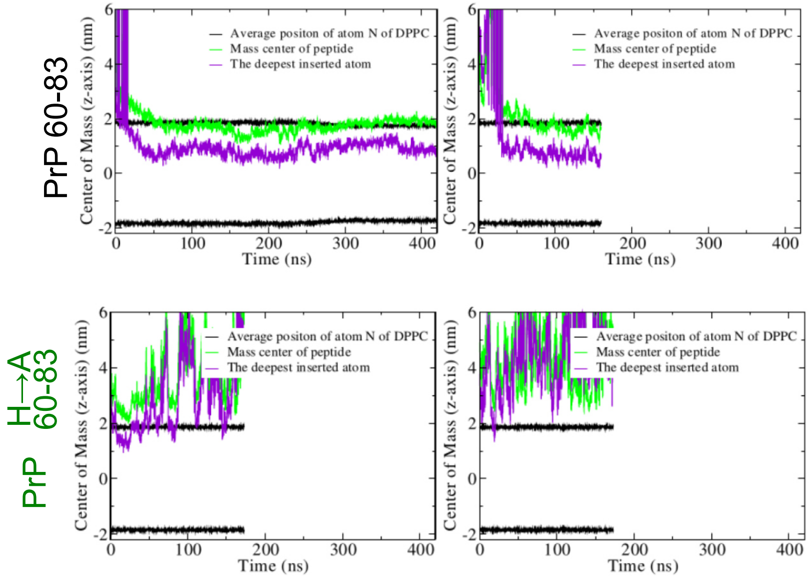


Reimann et al 2022, Supplement Figure 7

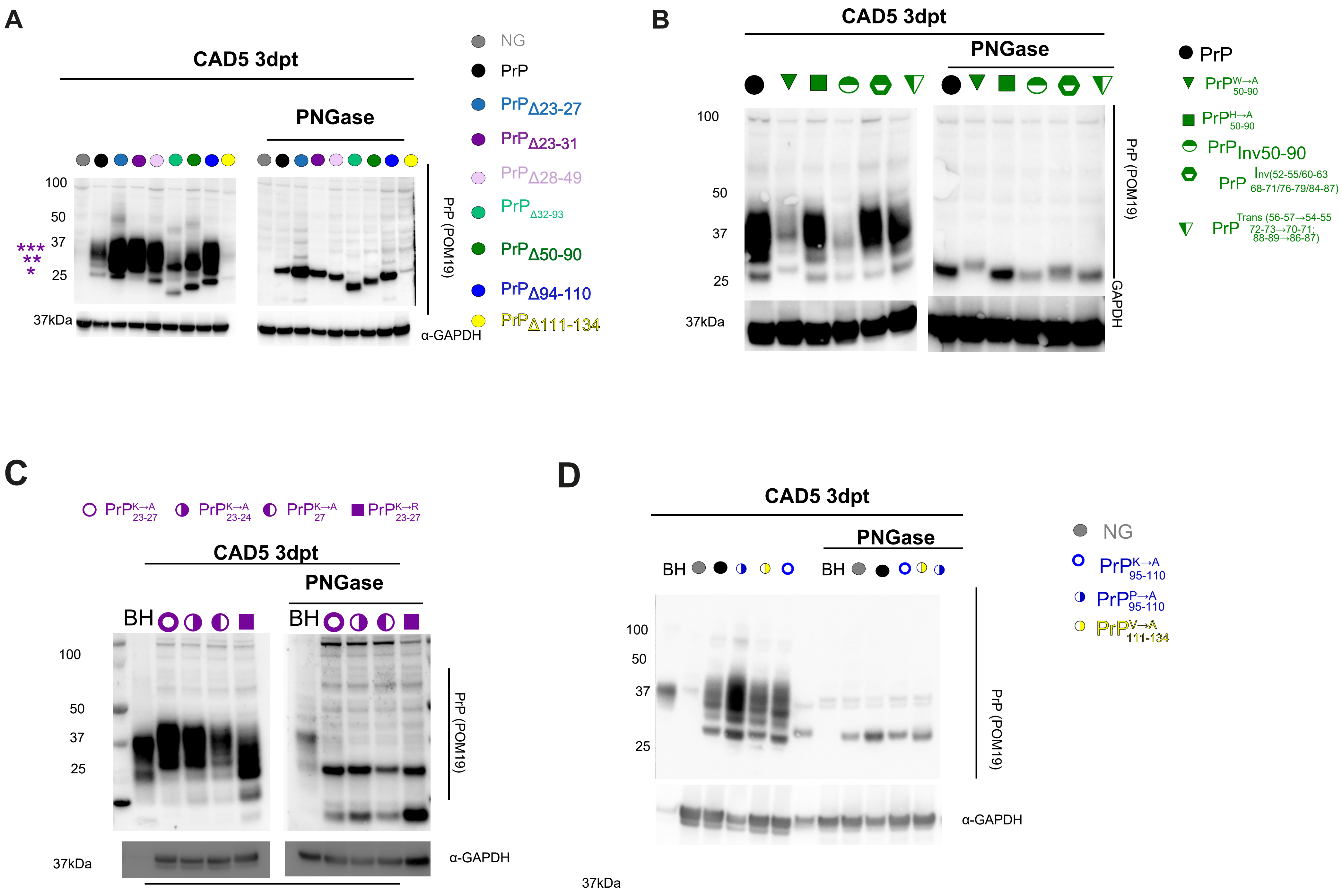


Reimann et al 2022, Supplement Figure 8

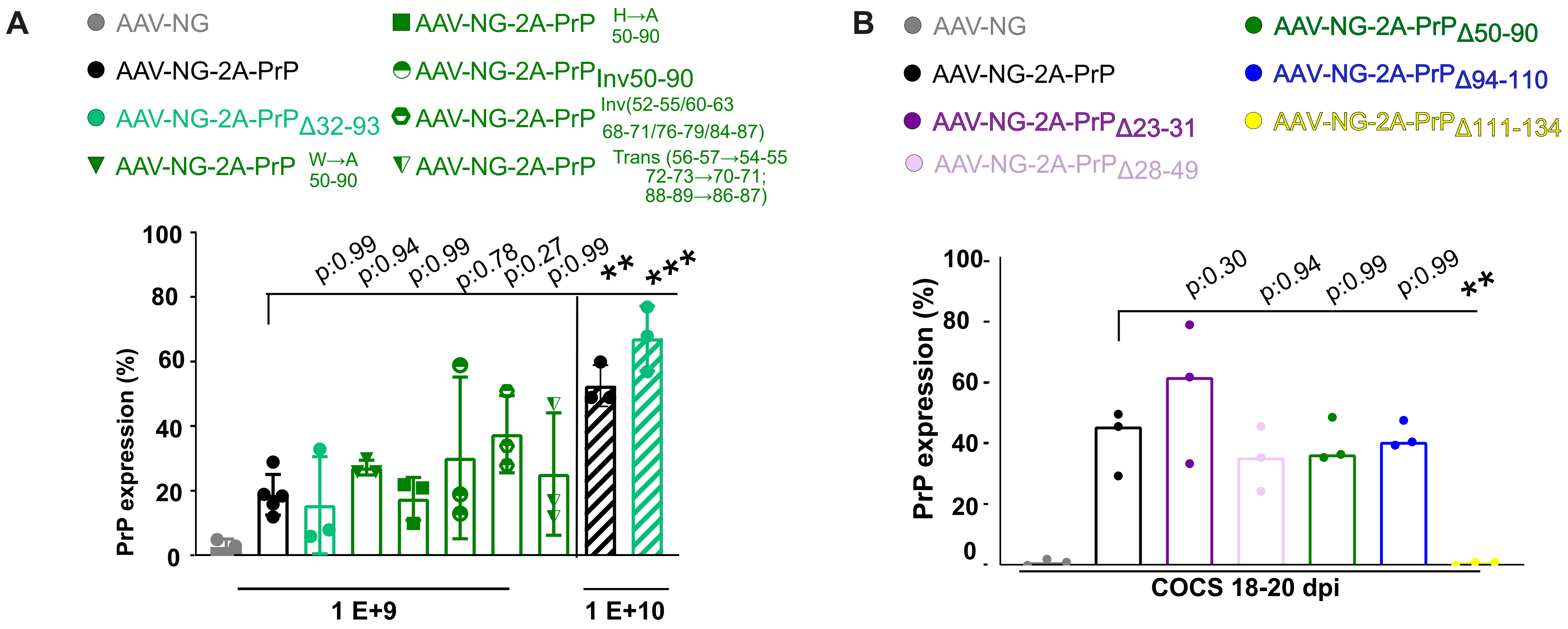


Reimann et al 2020, Supplement Figure 9
